# Multimodal brain imaging of insomnia, depression and anxiety symptoms: transdiagnostic commonalities and differences

**DOI:** 10.1101/2024.08.30.610439

**Authors:** Siemon C. de Lange, Elleke Tissink, Tom Bresser, Jeanne E. Savage, Danielle Posthuma, Martijn P. van den Heuvel, Eus J.W. van Someren

## Abstract

Insomnia disorder, major depressive disorder and anxiety disorders are the most common mental health conditions, with high comorbidity and genetic overlap suggesting shared brain mechanisms. Studies on brain correlates of these disorders have not fully addressed this overlap. Aiming to distinguish shared from specific brain structural and functional properties associated with symptoms of these disorders, this study analyzed multimodal brain imaging data from over 40,000 UK Biobank participants. Functional enrichment analyses were conducted to understand the cognitive-emotional and neurotransmission implications of the identified brain regions and connections. Results showed that smaller cortical surfaces, smaller thalamic volumes, and weaker functional connectivity were linked to more severe symptoms across all symptom types. Several symptom-specific associations were revealed, most commonly in different parts of the amygdala-hippocampal-medial prefrontal circuit. These findings revealed both transdiagnostically shared and unique brain properties that could lead to more directed treatment targets for insomnia, depression, and anxiety.

## Introduction

The three categories of mental disorders with the highest 12-month prevalence rates are insomnia, unipolar depression and anxiety (e.g. 7%, 6% and 14%, respectively, ^1^). These conditions have a large burden on the individual and prognosis is poor, due to moderate treatment success and a high chance of chronicity or relapse ^2–4^. Despite the urgent need for innovative transdiagnostic and disorder-specific treatments, targetable key mechanisms remain elusive.

Brain imaging studies have offered valuable biological insights into insomnia, depression and anxiety disorders ^5–7^. Yet, their intertwined nature – characterized by shared genetic risk factors, overlapping symptoms, high comorbidity, and causal interrelations ^8–11^ – underscores the urgency to explore their interconnected brain mechanisms more comprehensively. Although several studies have simultaneously addressed the brain correlates of anxiety and depression ^12–14^, most omitted a possible link with the highly comorbid insomnia. The disturbed sleep observed in insomnia disorder is hypothesized to be key to the development and severity of depression and anxiety by interfering with overnight relief of emotional distress ^2^. While a few studies have investigated the links of insomnia with either depression ^15–17^ or anxiety ^18–20^, these studies have often small sample sizes and report inconsistent results ^2^. Examining the three disorders together in a large-scale transdiagnostic study could provide insights into overlapping and distinct factors involved in insomnia, depression and anxiety.

In this study, we examined the brain structural and functional correlates of characteristics of anxiety, insomnia and depression using data obtained in over 40,000 participants from the UK Biobank. This approach allowed us to address a key question: can we distinguish brain property correlates that are unique to symptoms of a disorder from those that are shared across disorders? The UK Biobank dataset includes multimodal magnetic resonance imaging (MRI) data including brain measures such as cortical surface area, cortical thickness, subcortical volume, structural connectivity, functional connectivity and amygdala reactivity, enabling the integration of information from different modalities to obtain a comprehensive view of overlap and differences in the brain circuits associated with the severity of symptoms characterizing the three disorders.

## Results

We examined MRI and survey data of 41,667 participants from the UK Biobank. After (1) setting aside data from 5,000 participants for holdout validation, (2) discarding data from 4,893 participants due to incomplete MRI data or missing behavioral data, and (3) excluding 6,170 participants following data quality control, we had a discovery sample of 25,604 individuals (see Supplementary Methods and Supplementary Figure 1 for details). This sample comprised 53% female participants, and the participants had a median age of 64 years, spanning an age range from 45 to 81 years. Insomnia, depression and anxiety symptom severity scores were derived from UK Biobank self-report questionnaire data and showed small to moderate correlations (Spearman’s rank correlation coefficients: insomnia-depression *ρ*=0.18, p<0.001; depression-anxiety *ρ*=0.30, p<0.001; anxiety-insomnia *ρ*=0.16, p<0.001, all FDR-corrected).

We tested associations between brain measures and symptom severity scores using linear regression with age, gender, age-gender interactions and other methodological factors as covariates. Across six MRI modalities, we investigated differences in both global and regional brain metrics (see Supplementary Table 1 for an overview of all associations). Findings were followed up by functional enrichment analyses to discern potential cognitive-emotional, functional (presented in the Supplementary Results) and neurotransmission consequences arising from spatially distributed yet functionally linked brain association patterns.

### Cortical Surface Area

Individuals with a smaller *total* cortical surface area (CSA) exhibited more pronounced symptoms of insomnia, depression, and anxiety (β=-0.028, p<0.001; β=-0.032, p<0.001; and β=-0.050, p<0.001, respectively). While the association seemed most pronounced for the severity of symptoms of anxiety, differences between the strengths of associations did not reach significance (insomnia-depression: p=0.743; depression-anxiety: p=0.051; insomnia-anxiety: p=0.051, FDR-corrected).

*Regional* analyses pinpointed smaller CSA in 14, 35, and 61 regions in proportion to increasing severity of insomnia, depression, and anxiety symptoms, respectively, overlapping in the frontal regions (see Supplementary Table 1). Insomnia symptom severity was most strongly correlated with smaller CSA of the precentral gyrus (see Figure 1). The association of depressive symptom severity with smaller CSA was most evident in the frontal, middle and inferior temporal and parietal lobes. Anxiety symptom severity was most strongly correlated with smaller CSA in the insula, orbitofrontal and temporal regions.

**Figure 1.**
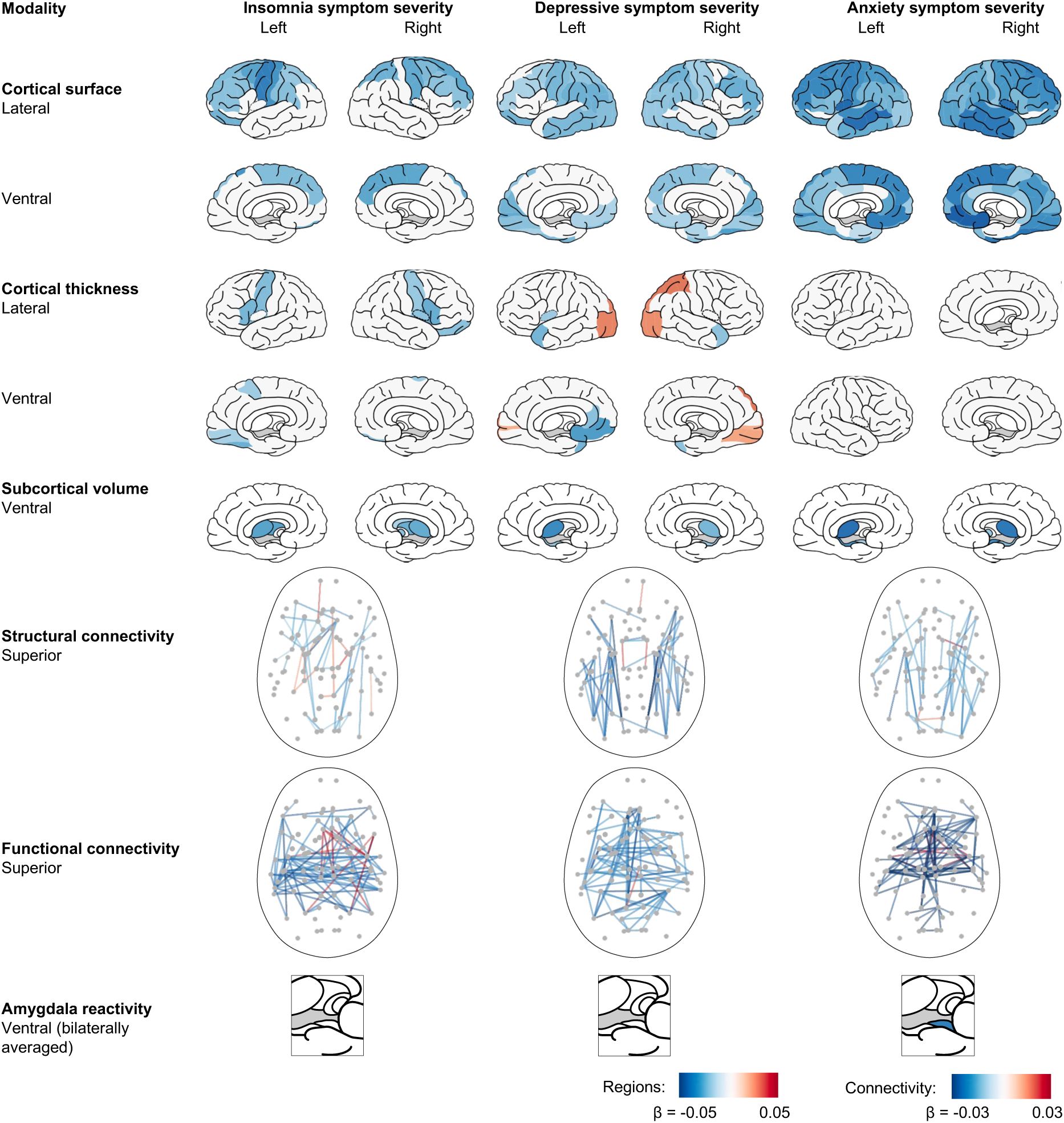
Associations of brain morphology and severity of insomnia, depression and anxiety symptoms. The first five rows present standardized regression coefficients for brain areas whose surface areas, thicknesses, or volumes are significantly associated with the severity of symptoms of insomnia, depression, or anxiety (p<0.05, FDR-corrected across 68 tests for cortical surface and thickness, and 14 tests for subcortical volume). The sixth and seventh rows show the top 10% structural and functional connections with the highest absolute (positive or negative) association strengths. The eighth-row highlights that bilaterally averaged amygdala reactivity to angry or fearful faces is only correlated with anxiety symptom severity (p<0.05, FDR-corrected across three tests). Negative associations (in blue) indicate smaller surface, thickness and volume, weaker structural and functional connectivity, and lower amygdala reactivity in people with more severe symptoms.

*Functional* annotation showed that cortical regions with smaller CSA in association with the severity of insomnia, anxiety and depressive symptoms overlapped in five of the eight cognitive-emotional domains and 13 of the 18 neurotransmission systems (see Figure 2 and Table 1 and 2 for a summary of the findings and Supplementary Table 2 and 3 for an overview of all associations). More specific to the severity of insomnia than depressive symptoms were smaller CSAs in areas with high expression of alpha-4 beta-2 nicotinic (α4β2) receptor, histamine H3 receptor, and norepinephrine (NET) transporter (p=0.002, p=0.041, and p=0.047, respectively, FDR-corrected). More specific to the severity of depressive than insomnia symptoms were smaller CSAs in regions tied to the functional domain of “vision” (p=0.037, FDR-corrected) and those enriched with dopamine D2 receptors (p=0.021, FDR-corrected). More specific to the severity of anxiety than depressive symptoms were smaller CSAs in regions implicated in the “reward” domain (p=0.037) and regions with high densities of the cannabinoid receptor type 1 (CB1), dopamine D1, histamine H3, and MOR receptors (p=0.002, p=0.019, p=0.006, and p=0.046, respectively). More specific to the severity of anxiety than insomnia symptoms were smaller CSAs in regions enriched for M1, D2 and 5HT1A receptors (p=0.006, p=0.002 and p=0.041, respectively).

**Figure 2.**
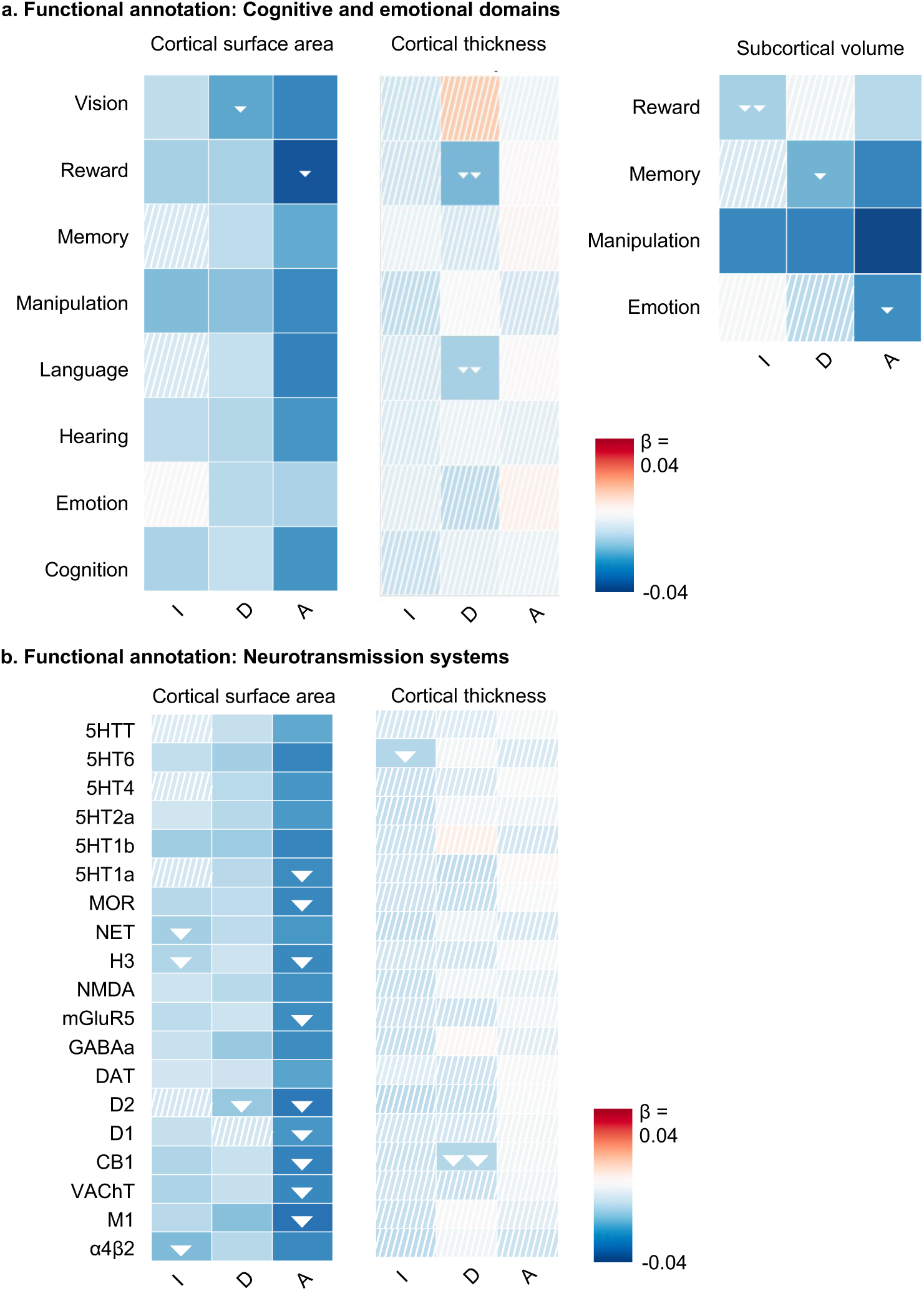
Functional annotation of regional brain associations. Enrichment analysis evaluated whether regional symptom-related variations were enriched within any cognitive-emotional or neurotransmission system. **a.** These correlation maps show the associations between symptom severities and the cortical surface area, thickness and subcortical volume in regions linked to cognitive-emotional domains. Associations that are statistically significant (p<0.05 after FDR-correction across 24 tests for cortical surface and cortical thickness [3 comparisons x 8 domains], and 12 tests for subcortical volume [3 comparisons x 4 domains]) are depicted in solid color, while non-significant associations are marked with hatching. Negative effects (in blue) denote that the involved regions had smaller surface area, thickness or volume in specific cognitive-emotional domains or neurotransmission systems. Additionally, white arrows highlight that, compared to another symptom, the association of the severity of a symptom is significantly more aggregated in areas involved in the indicated domain. **b.** Correlation maps that present the associations of symptom severities with the cortical surface area and cortical thickness of the top 25% cortical regions that have the highest densities for each of 18 receptor and transporter types (p-values are FDR-corrected across 54 tests [3 comparisons x 18 receptors and transporters]).

**Table 1.** Overview of significant findings for each cognitive-emotional domain across modalities. In general, as the severity of symptoms of insomnia (I), depression (D) and/or anxiety (A) increases, people have smaller surface area, thinner cortex, or less structural or functional connectivity between brain regions involved in the cognitive-emotional domain, except for two associations (indicated in the table by an upward arrow ↑): For connections between regions in the emotional domain, more severe insomnia symptoms were associated with stronger structural connectivity; and for connections between regions in the language domain, more severe depressive symptoms were associated stronger structural. A single asterisk (*) denotes that the association between symptom severity and brain measure deviation is significantly more aggregated in connectivity or areas involved in the indicated domain compared to one other symptom type. Double asterisks (**) indicate that relative to both other symptom types, the symptom severity-brain measure association is more aggregated to areas or connectivity involved in the indicated domain.

|  | Cortical surface area | Cortical thickness | Subcortical volume | Structural connectivity | Functional connectivity |
| --- | --- | --- | --- | --- | --- |
| **Cognition** | I D A |  |  | I* D | I A* |
| **Emotion** | D A |  | A* | I↑ D A* | I A |
| **Hearing** | I D A |  |  | I* | A |
| **Language** | D A | D** |  | D↑ | I D A |
| **Manipulation** | I D A |  | I D A | D A | I D A |
| **Memory** | D A |  | D* A | D* | D A |
| **Reward** | I D A* | D** | I** A |  |  |
| **Vision** | I D* A |  |  | I D A | D |

**Table 2.** Overview of significant findings for each receptor or transporter across modalities. All significant associations are in the same direction: in proportion to the severity of symptoms of insomnia (I), depression (D) and/or anxiety (A), people have a smaller surface area, thinner cortex, or less structural or functional connectivity between areas with high expression of the receptor or transporter. A single asterisk (*) denotes that the association between symptom severity and brain measure deviation is significantly more aggregated in areas or connectivity involved in the indicated domain compared to one other symptom type. Double asterisks (**) indicate that relative to both other symptom types, the symptom severity-brain measure association is more aggregated in areas or connectivity involved in the indicated domain.

|  | Cortical<br>surface area | Cortical<br>thickness | Structural<br>connectivity | Functional<br>connectivity |
| --- | --- | --- | --- | --- |
| $\alpha 4\beta 2$ | I* D A | | I D | I* A* |
| M1 | I D A* |  | A | D A |
| VACHT | I D A* |  |  | D A |
| CB1 | I D A* | D** |  | I D A |
| D1 | I A* |  |  | I D A** |
| D2 | D* A* |  |  | I D A |
| DAT | I D A |  |  | I D A* |
| GABAa | I D A |  | D | D A |
| mGluR5 | I D A* |  |  | I D A** |
| NMDA | I D A |  |  | I D A** |
| H3 | I* D A* |  | I | I D A** |
| NET | I* D A |  | D | D A |
| MOR | I D A* |  | I | I D A* |
| 5HT1a | D A* |  | D | I D A |
| 5HT1b | I D A |  | I D A | I A |
| 5HT2a | I D A |  |  | I D A |
| 5HT4 | D A |  |  | I D A |
| 5HT6 | I D A | I* | I D A | I D A |
| 5HTT | D A |  | A | I D A |

### Cortical thickness

People showed a thinner *average* cortical thickness in proportion to the severity of insomnia (β=-0.015, p=0.027) but not depressive (β=-0.008, p=0.24) or anxiety symptoms (β=-0.005, p=0.45). The strength of the association with cortical thickness did however not differ significantly across symptom types (all p>0.05, FDR-corrected).

*Regional* analysis pinpointed thinner cortices in nine regions in proportion to insomnia severity, including the left fusiform area, bilateral pars opercularis and precentral gyrus (p<0.05, FDR-corrected, see Figure 1 and Supplementary Table 1). In proportion to depressive symptom severity, thinner cortices were found in eleven regions, including the left medial frontal region and the temporal lobe, and thicker cortices in occipital regions. Anxiety severity was not associated with regional cortical thickness.

*Functional* annotation showed that cortical areas that were thinner in association with insomnia severity did not map on any specific cognitive-emotional domain but did map on regions with high 5HT6 serotonin receptor densities (β=-0.012, p=0.040), and significantly more so for insomnia than for depressive symptoms (p=0.034, see Figure 2). Thinner cortices in proportion to depressive symptom severity occurred in regions related to the “reward” (β=-0.018, p=0.002) and “language” (β=-0.014, p=0.020) domains, and significantly more so than was the case for the severity of insomnia (p=0.006 and p=0.002) or anxiety symptoms (both p<0.001). Thinner cortices in proportion to depressive symptom severity also mapped on regions with higher cannabinoid CB1 receptor density (β=-0.012, p=0.040), and significantly more so than was the case for the severity of insomnia (p=0.001) and anxiety (p<0.001).

### Subcortical volume

A smaller *total* subcortical volume was linked to more severe symptoms of insomnia (β=-0.019, p=0.011) and anxiety (β=-0.031, p<0.001), but not depression (β=-0.014, p=0.061). No significant symptom type differences were found when comparing the severity association strengths (all p>0.05, FDR-corrected).

*Regional* analyses showed that severity of all three symptom types was associated with smaller thalamic volume bilaterally (see Figure 1 and Supplementary Table 1). Furthermore, in proportion to insomnia symptom severity, the bilateral pallidum, right nucleus accumbens, and right caudate nucleus showed smaller volumes. More severe depressive symptoms were found in people with smaller hippocampal volumes bilaterally, and more severe anxiety symptoms in people with smaller volumes of the hippocampus and amygdala bilaterally, and of the right nucleus accumbens.

*Functional* annotation showed that smaller subcortical volumes in proportion to the severity of insomnia symptoms occurred in the “manipulation” (β=-0.026, p<0.001) and “reward” (β=-0.014, p=0.014) domains. Mapping on the “reward” domain was significantly more pronounced for insomnia severity than for the severity of anxiety (p=0.003) or depressive symptoms (p<0.001). Smaller volumes in proportion to depressive symptoms severity occurred in regions associated with the “manipulation” (β=-0.027, p<0.001) and “memory” (β=-0.019, p=0.008) domains. Mapping on the “memory” domain was significantly more pronounced for depressive symptom severity than for insomnia symptom severity (p=0.027). Smaller volumes in proportion to anxiety symptom severity occurred in regions related to the “manipulation” (β=-0.037, p<0.001), “memory” (β=-0.027, p<0.001), “reward” (β=-0.012, p=0.041) and “emotion” (β=-0.025, p<0.001) domains. Mapping to the “emotion” domain was significantly more pronounced for anxiety symptom severity than for insomnia symptom severity (p<0.001).

### Structural connectivity

The *average* structural connectivity strength, measured by fractional anisotropy, was inversely related to the severity of depressive (β=-0.019, p=0.003) and anxiety symptoms (β=-0.016, p=0.014), but not to insomnia symptom severity (β=-0.006, p=0.37). The differences in association strength did however not reach significance (all p>0.05, FDR-corrected).

At the *regional* level, insomnia symptom severity was associated with the connectivity from five brain regions; depressive symptom severity with the connectivity of 42 regions; and anxiety symptom severity with the connectivity of 38 regions (see Figure 3a and Supplementary Table 1). Insomnia symptom severity was associated with the strength of connectivity of regions including the superior frontal and lingual gyrus and cuneus. Depressive symptom severity was most notably associated with connectivity strength of the inferior parietal gyrus, lateral occipital gyrus, and pericalcarine gyrus. Anxiety symptom severity was most notably associated with connectivity strength of the lateral occipital gyrus, hippocampus, and cuneus.

**Figure 3.**
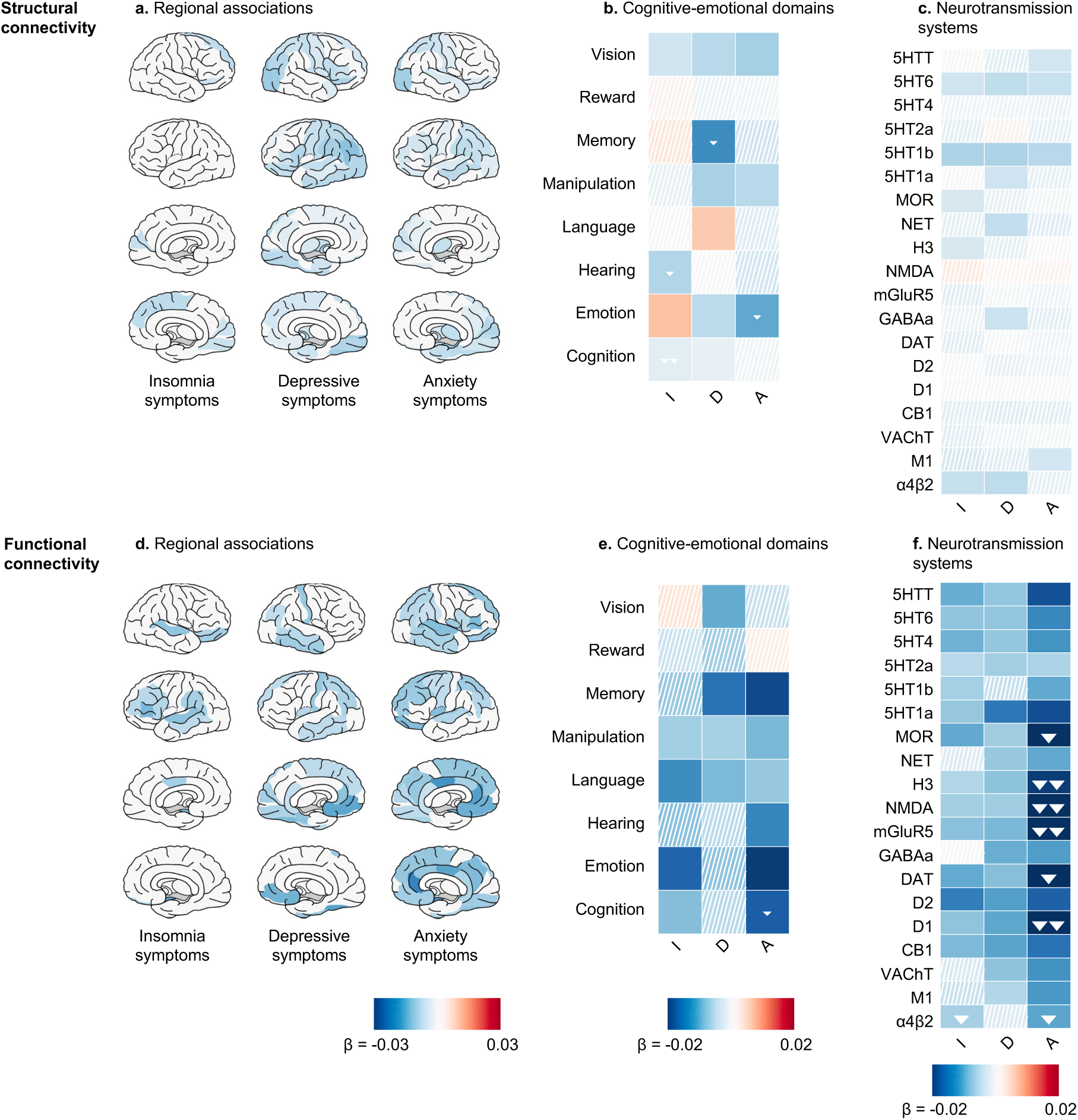
Associations of structural and functional connectivity with symptom severity of insomnia, depression and anxiety. **a.** and **d.** Connectivity deviations of brain regions. Brain areas with region-wise connectivity that deviated significantly in proportion to symptom severity are colored (p<0.05, FDR-corrected for 82 tests). Negative effects (in blue) indicate weaker connectivity of the regions in people that experienced more severe symptoms. **b.** and **e.** Associations of symptom severities with deviations in connectivity strength between areas involved in 8 cognitive-emotional domains. Significant associations (p<0.05 after FDR-correction across 24 tests [3 comparisons x 8 domains]) are in solid color, non-significant associations are hatched. **c.** and **f.** The associations of the functional connectivity of 18 neurotransmission system with the three symptom types (p-values being FDR-corrected across 54 tests [3 comparisons x 18 receptors and transporters]). In panels **b. c. e.** and **f.**, white arrows indicate that, compared to one other symptom type, the association of the severity of a symptom with the degree of deviation in connectivity strength is significantly more pronounced in areas involved in the indicated domain. Cells featuring two white arrows indicate that, relative to both other symptom types, the symptom severity-connectivity associations were more concentrated in areas involved in the indicated domain.

*Functional* annotation analyses highlighted that symptom-related connectivity strength clustered in various cognitive-emotional domains and neurotransmission circuits (see Figure 3b and 3c). Across the three symptom types, more severe symptoms were associated with weaker connectivity in ‘vision’-related regions and regions with high 5HT6 serotonin receptor density. Weaker connectivity specifically in proportion to the severity of symptoms of insomnia but not depression occurred for areas implicated in the ‘hearing’ domain (p=0.027). Moreover, weaker connectivity specifically in proportion to the severity of symptoms of insomnia but not depression or anxiety occurred for areas implicated in the ‘cognition’ domain (both p=0.027). Weaker connectivity specifically in proportion to the severity of symptoms of depression but not insomnia occurred for areas implicated in the ‘memory’ domain (p=0.027). Lastly, weaker connectivity specifically in proportion to the severity of symptoms of anxiety but not insomnia occurred for areas implicated in the ‘emotion’ domain (p=0.014).

### Functional connectivity

Individuals with weaker *average* functional connectivity exhibited more pronounced symptoms of insomnia, depression, and anxiety (β=-0.013, p=0.037; β=-0.018, p=0.007; β=-0.023, p<0.001, respectively). The strength of these associations did not differ for tree types of symptoms (all pairwise p>0.05, FDR-corrected).

At the *regional* level, more severe insomnia symptoms were associated with weaker connectivity of 14 brain regions, notably of the bilateral amygdala and superior temporal regions (see Figure 3d and Supplementary Table 1). More severe depressive symptoms were associated with weaker connectivity of 20 regions, including the fusiform and medial orbitofrontal regions. More severe anxiety symptoms were associated with weaker connectivity in 43 regions, particularly of the cingulate cortex and amygdala.

*Functional* annotation analyses showed an overlap across the three symptom type severities in the mapping of connectivity deviations to the ‘manipulation’ and ‘language’ domains and to 13 neurotransmission systems (see Figures 3e and 3f). More specific to the severity of insomnia symptoms than depressive symptoms was weaker connectivity between regions with enriched α4β2 receptor densities (p=0.003). The 20 regions with weaker connectivity in proportion to the severity of depressive symptoms did not map to specific functional domains. More specific to the severity of symptoms of anxiety as compared to depression or insomnia was the weaker connectivity between regions with high densities of D1 (p=0.001 and p=0.008), H3 (p=0.003 and p=0.006), and mGluR5 (p=0.006 and p=0.008) receptors. Furthermore, more specific to the severity of symptoms of anxiety as compared to depression was the weaker connectivity of regions in the “cognition” domain (p<0.001) and of regions with high densities of MOR (p=0.001) and α4β2 (p=0.003) receptors and dopamine transporter (DAT, p=0.003).

### Amygdala responsivity

Task-based fMRI showed that amygdala reactivity to angry or fearful face stimuli was inversely related to the severity of anxiety symptoms (β=-0.034, p<0.001). Such association was not observed for insomnia (β=-0.013, p=0.270) or depressive symptoms (β=-0.007, p=0.073, FDR-corrected). Comparative analyses showed a stronger association of amygdala responsiveness with anxiety symptom severity than with insomnia (p=0.021) or depressive symptom severity (p=0.001, FDR-corrected).

### Potential confounding factors

We assessed the impact of potential confounding factors by including them as covariates in the association analyses of the regional and connectivity brain properties. The inclusion of socio-economic status, educational attainment, smoking, alcohol use, diabetes, and hypertension did not significantly alter the results (refer to Supplementary Figure 2). Indicators of chronic distress, such as the number of traumatic experiences during childhood and adulthood, along with psychiatric medication use (including anxiolytics and antidepressants) reduced the number of replicated regional associations significantly for depressive and anxiety symptoms, but not for insomnia symptoms (see Supplementary Table 4). We also observed correlations between the symptom scores and these covariates, suggesting collinearity (see Supplementary Figure 3). Including body mass index (BMI) as covariate significantly altered the regional associations of cortical thickness with depression severity, although it had no significant impact on the other associations.

### Sensitivity and validation analyses

To evaluate the robustness and generalizability of our findings, we conducted a series of sensitivity and validation analyses. Results obtained from sensitivity analyses applying a different atlas (Desikan-Killiany DK-114), alternative symptom severity scores and applying statistical testing using spatial spin-permutation based null-models are presented in the Supplementary Results. Further, a sensitivity analysis that excluded 3,532 participants with familial ties to evaluate possible genetic confounding, showed no significant effect on the results (see Supplementary Results).

*In-sample validation* analyses utilized MRI and behavioral data from a second MRI session conducted 2-3 years later with 2,309 participants. *Out-of-sample validation* analyses utilized the holdout sample of 4,076 participants. Detailed results are available in the Supplementary Results. Consistent across all validation analyses were the smaller total cortical surface area, weaker average functional connectivity and weaker amygdala reactivity in participants with more severe anxiety symptoms (see Supplementary Table 5). At the regional level, findings from the analyses aligned with the initial findings for the regional associations of insomnia symptoms with cortical thickness, anxiety symptoms with cortical thickness, subcortical volume and functional connectivity, and depressive symptoms with subcortical volume (see Supplementary Figures 4 and 5).

## Discussion

This study revealed overlap between and differences in the multimodal brain imaging fingerprints of symptoms related to insomnia, depression, and anxiety disorders in participants from the UK Biobank. We identified shared and specific neurobiological correlates of symptom severities. Symptom-specific correlates suggested differential deviations converging in the amygdala-hippocampus-medial prefrontal cortex circuit.

Individual variability in the severity of all three symptom types correlated with several brain properties. Globally, people with more severe symptoms had a smaller total cortical surface area and weaker average functional connectivity. Regionally, all symptom severities were inversely associated with thalamic volume. These findings are consistent with previous studies reporting a smaller thalamic volume in insomnia, depression, and anxiety ^7,21–23^. It is possible that the shared associations extend beyond the severity of insomnia, depression and anxiety, and hold as well for other psychiatric conditions ^24,25^.

Our findings also shed light on the symptom-specific nature of symptoms observed in earlier single-disease studies. The link of weaker structural connectivity in “cognition”-related areas specifically with more severe insomnia symptoms is in line with an earlier study reporting reduced fronto-subcortical connections in insomnia ^26^ and supports the disorder-specific nature. The thinner gray matter within the “reward” and “language” domains and within the “limbic” functional network specifically in people with more severe depressive symptoms, aligns with previous studies that reported thinner cortices in the temporal pole and medial orbitofrontal cortex in depressive patients ^6^. Lower amygdala reactivity to fearful and angry faces was specifically associated with higher anxiety symptom severity. Two recent studies also did not find altered amygdala reactivity in association with either depression or insomnia ^27,28^. Certain associations of brain features with symptom severity appeared also specific to two out of three symptom types (see Figure 4). Brain areas and circuits with such associations mapped onto the dopamine, histamine and nicotinic neurotransmission systems and are further discussed in the Supplementary Discussion.

**Figure 4.**
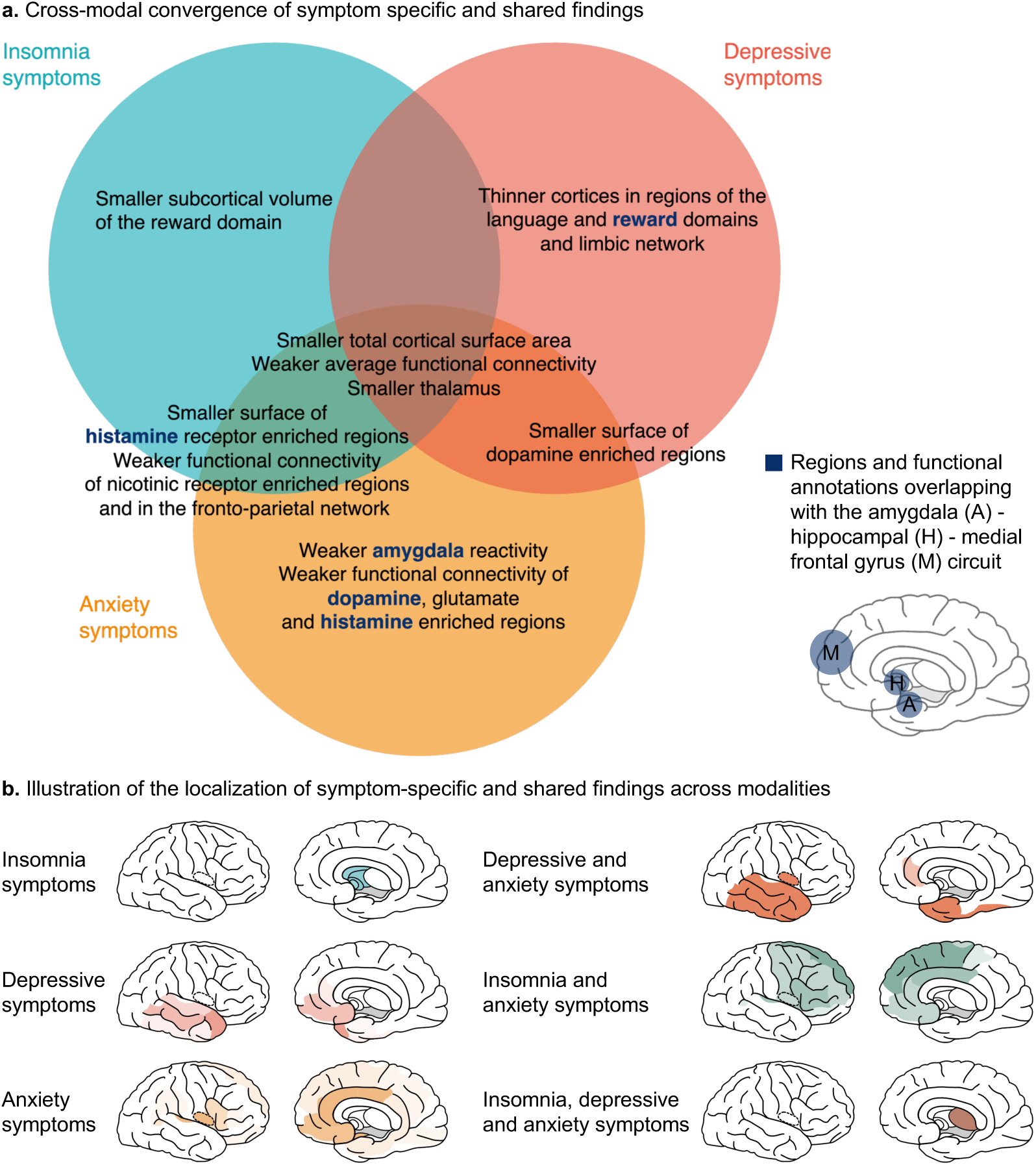
Summary of Cross-Modal Findings. **a.** Comparative analysis showed that specific brain properties, cognitive-emotional domains, and neurotransmission systems are linked to the severity of certain symptoms, or combinations of them. Across symptom-specific and shared findings, we see regions and systems that overlap with the amygdala-hippocampal-medial prefrontal circuit (highlighted in blue and bold text). Known for its role in stress-related processes and REM sleep, the involvement of this circuit in all three symptoms supports the hypothesis that it plays a central role in their shared etiology. **b.** Illustration of the spatial localization of the regions, cognitive-emotional domains and neurotransmission systems associated with specific symptoms or symptom combinations.

The multimodal analyses provided insights into how brain areas and circuits associated with symptom severity can present across imaging modalities, discussed further in the Supplementary Discussion. We noted a cross-modal convergence of areas and circuits of which features were associated with the severity of both insomnia and anxiety. Insights in cross-modal convergence and divergence offer a relational perspective on potential brain mechanisms, which could be further investigated to construct transmodal biomarkers ^29^.

Reviewing the findings from this study highlights that the brain correlates specific to one or two symptoms predominantly occur in the amygdala-hippocampus-medial prefrontal cortex (MPC) circuit (see Figure 4). This convergence occurs across modalities and hints at a shared neurobiological circuit key to the vulnerability of anxiety, depression, or insomnia. This circuit has been implicated in stress-related processes ^30^ and in REM sleep ^31^, which have both been reported to deviate in the three disorders. This circuit may be an important substrate for altered overnight plasticity with restless REM sleep, proposed to underly insufficient overnight resolution of emotional distress ^2,32^. Thus, while our study found, in line with previous studies ^33^, only weak associations between brain measures and symptom severity (|β|<0.05), where a one standard deviation change in a brain measure was associated with a maximum 1.10-fold higher risk of experiencing symptoms, our population-based findings may direct focus for future translational patient-control imaging studies with in-depth phenotyping. Focus for therapeutic drug development is moreover provided by functionally annotating involved areas and circuits to neurotransmission systems (see Supplementary Discussion).

In interpreting our findings, several inherent limitations should be considered. First, symptom severity scores do not map one-to-one to diagnosed disorders and can only be considered proxy measures of the general characteristics of these disorders. While these scores are grounded in literature and validated (see Supplementary Methods), limitations are inherent to the use of a small set (1-3) of questions to measure the severity of symptoms of complex disorders. Furthermore, it is important to interpret these scores within the broader context of their interconnected relationships with other variables, such as past exposure to distress and the use of antidepressants and anxiolytics, which showed to overlap with the symptom severity scores (see Supplementary Figure 2). For insomnia severity, averaging scores from two time points, our study may have been geared more toward symptom severity in chronic insomnia than acute insomnia, possibly contributing to differences with studies that utilized a single time point ^5,34^. A second limitation is that the UK Biobank sample is older, more educated, and healthier than the general UK population ^35^. A third limitation, inherent to the small effect sizes of the reported associations, is that replication would require a very large dataset with a similar participant age range as the UK Biobank. Future replication studies could be performed when such a dataset becomes available. A final limitation to be mentioned is the variation in power to detect associations across different regions of the brain, connections, and cognitive domains ^36^. Methodological factors can influence the ability to detect changes in a specific brain region or modality. Consequently, the absence of significant effects in certain regions or systems does not necessarily imply an absence of deviations. Future studies with increased power may find that some associations reported here as symptom-specific, are also in part common across symptoms.

In conclusion, our findings underscore the importance of an integrative approach to understand insomnia, depression, and anxiety. The findings moreover revealed the transdiagnostic importance of the amygdala-hippocampus-medial prefrontal cortex circuit. These findings are a starting point for further research into these interconnected disorders to test targeted hypotheses in clinical settings.

## Methods

### Sample

This study used data from the UK Biobank database, a population-based sample of adults in the UK who volunteered to complete an in-person visit and link their data from the national health registry ^37^. The study was conducted under application number 16406. All participants provided written informed consent, and the UK Biobank received ethical approval from the National Research Ethics Service Committee North West–Haydock (reference 11/NW/0382) and all procedures were conducted following the World Medical Association declaration of Helsinki. Within the UK Biobank database, demographics and health-related data from questionnaires and electronic records were available for N=502,537 participants in March 2022 (see Supplementary Figure 1). Neuroimaging data were available for a subsample of N=41,667 individuals, of which N=3,601 individuals had also neuroimaging data available from a second (repeat) visit that was used in a replication analysis (release January 2020). Before any analyses for this study were conducted, we randomly assigned 5,000 participants to a holdout set that was used for validation, resulting in a research set of N=36,667, which underwent a subsequent quality control procedure.

### Image processing

Multimodal analyses were performed based on pre-processed T1-weighted MRI, T2-flair weighted MRI, resting-state functional MRI (rs-fMRI), task-based functional (t-fMRI) and diffusion weighted imaging (DWI) provided by the UK Biobank. The UK Biobank scanning protocols and (pre)processing pipeline are described by Miller and colleagues ^38^ and in the UK Biobank Brain Imaging Documentation (https://biobank.ctsu.ox.ac.uk/crystal/crystal/docs/brain_mri.pdf).

#### Structural brain measures

The UK Biobank provides cortical surface reconstructions and brain segmentations (field ID: 20263) obtained using FreeSurfer (v6.0) ^39^ that were based on T1 and T2-flair weighted MRI data. From the FreeSurfer output, the cortical thickness and cortical surface area of the 68 brain regions in FreeSurfer’s Desikan-Killiany atlas were computed ^40^. Furthermore, the volumes of 14 subcortical brain regions were obtained from FreeSurfer’s automatic subcortical segmentation.

#### Structural connectivity

Structural connectivity was reconstructed using Connectivity Analysis Toolbox (CATO, v3.1.1) ^41^. The UK Biobank provided preprocessed DWI files that were corrected for eddy currents, head motion and outlier-slices and gradient distortions (field ID: 20250). Parcellation was applied to the DWI data using an affine transformation mapping the T1-weighted image to the diffusion-weighted image ^42^. The white matter fiber organization in each voxel was estimated using super-resolved constrained spherical deconvolution (spherical harmonics order=6, *λ*=1, *τ*=0.1, number of output peaks=4, minimum peak ratio=0.1, maximum number of peaks=8) ^43^. White matter fibers were reconstructed by the Fiber Assignment by Continuous Tracking (FACT) algorithm ^44^, with streamline reconstruction starting from 8 seeds in all white matter voxels. Fiber tracking was continued until a streamline showed high curvature, exited the brain mask or when a streamline entered a voxel with low fractional anisotropy (FA<0.1). Fractional anisotropy was estimated from the diffusion tensor that was measured in a voxel using the informed Robust Estimation of Tensors by Outlier Rejection (iRESTORE) algorithm ^45^ with automated parameter selection ^46^. The mean fractional anisotropy (FA) of a streamline was computed as the weighted average over all voxels that a streamline passed.

#### Functional connectivity

Functional connectivity was reconstructed using CATO (v3.1.1) ^41^. The UK biobank provided preprocessed fMRI data that were motion corrected, intensity normalized, high-pass filtered and corrected for echo-planar imaging (EPI) and gradient distortions, and artifacts were removed using FMRIB’s independent component analysis (ICA) based X-noiseifier (ICA+FIX, field ID: 20227). Using an affine transformation mapping the T1-weighted image to the rs-fMRI data, the FreeSurfer parcellation was applied to the rs-fMRI data ^42^. Per voxel, the rs-fMRI time series was corrected for covariates including the linear trend and first order drift of the motion parameters, mean signal intensity of voxels in the white matter and corpus callosum. The time series were band-pass filtered between 0.01 – 0.1 Hz. Frames displaying significant motion artifacts, and one frame prior to the labelled motion artifacts, were removed from the rs-fMRI time series ^47^. Motion artifacts were defined as frames with either framewise displacement (FD) larger than 0.25 or a change in signal intensity between frames (DVARS) larger than Q3 + 1.5 × IQR, where IQR refers to the interquartile range (IQR=Q3 – Q1) and Q1 and Q3 refer to the first and third quartiles of the DVARS of all frames, respectively. Region-to-region functional connectivity was quantified by Pearson’s correlation coefficient of the average signal intensity of the voxels in the regions.

#### Amygdala responsivity

Amygdala responsivity to emotional faces is an imaging-derived phenotype provided by the UK Biobank (field ID: 25052). This measure was derived from a 4-minute task-based fMRI scan, during which participants performed the Hariri face-matching task ^48^. In this task, blocks of shapes or emotional faces (with angry or fearful expressions) are presented on a screen and participants are instructed to match the target shape or face from the top of the screen to one of the two options at the bottom. The faces-shapes contrast was computed to measure the amygdala response to emotional faces corrected for a general visual response to shapes. Amygdala responsivity was calculated as the median BOLD (blood-oxygen-level dependent) effect for the faces-shapes contrast in a group-defined amygdala responsivity mask.

### Quality control

A strictly data-driven quality control procedure was applied to the imaging data of discovery sample, second imaging visit and holdout validation sample (described below). For the discovery sample, participants with unsuccessful MRI data, as indicated by the UK Biobank (“unusable” data ^49^) or errors in the processing pipeline, were excluded, resulting in successfully processed imaging data of N=31,827 subjects. Secondly, a data driven quality control was performed based on the quality assessments derived from the raw and derived MRI data resulting in UK Biobank provided measures and data-based measures resulted in a sample size of N=25,657 (See Supplementary Methods).

### Symptom severity measures

Insomnia, depression and anxiety symptoms were measured using UK Biobank self-report questionnaire data that were collected at three time points: the initial assessment center visit, the imaging visit and the repeated imaging visit. The symptom scores averaged over the initial assessment visit and imaging visit were used in the main analyses with the aim to capture chronic rather than incidental symptom severity. Symptom scores from the repeated imaging visit were used in the replication analyses. A detailed validation of the symptom severity scores against diagnostic data and other in-depth questionnaires and an alternative severity score for anxiety and depressive symptoms, used in the sensitivity analyses, are presented in the Supplementary Materials.

Insomnia symptom severity was assessed by means of the question “Do you have trouble falling asleep at night or do you wake up in the middle of the night?” (field ID: 1200) and responses “never/rarely” rated 0, “sometimes” rated 1, “usually”, rated 2^50^. Participants that answered “Do not know” or “Prefer not to answer”, or for which no data was available, were excluded. Averaging over two assessments resulted in an insomnia severity score range from 0 to 2 in steps of 0.5.

Depressive symptoms were assessed by two questions from the PHQ-2 ^51^ questionnaire: “Over the past two weeks, how often have you felt down, depressed or hopeless?” (field ID: 2050) and “Over the past two weeks, how often have you had little interest or pleasure in doing things?” (field ID: 2060), scoring each as 0 for “not at all”, 1 for “several days”, 2 for “more than half the days” or 3 for “nearly every day”. Averaging the two PHQ-2 questions over two assessments resulted in a depression severity score range from 0 to 6 in steps of 0.5.

Anxiety symptoms were measured by three questions that correspond to the salient items from the anxious-tense factor ^52,53^ identified in the Short Scale of the Eysenck Personality Questionnaire-Revised (N-12) ^54^: “Would you call yourself a nervous person?” (field ID: 1970), “Would you call yourself tense or ‘highly strung’?” (field ID: 1990) and “Do you suffer from ‘nerves’?” (field ID: 2010). Affirmative answers (Yes) were scored as 1, negative answers (No) were scored as 0 and the anxiety symptom level was calculated as the sum of the three questions. Averaging over two assessments resulted in a depression severity score range from 0 to 3 in steps of 0.5.

### Analysis

Associations of severity of symptoms with structural and functional brain measures were estimated using linear regression models. Separate models estimated the predictive value of each brain measure (total, regional or connectivity measure) for either insomnia, depression or anxiety symptom severity as outcome variable. Follow-up analysis (see below) tested for symptom-specificity of associations. Models were adjusted for common covariates ^55^ including age (field ID: 21003), sex (field ID: 31), scanner site (field ID: 54), height (field ID: 12144), age*age, age*sex, handedness (field ID: 1707), scanner lateral (X) brain position (field ID: 25756), scanner transverse (Y) brain position (field ID: 25757), scanner longitudinal (Z) brain position (field ID: 25758), scanner table position (field ID: 25759) and intensity scaling of T2 FLAIR (field ID: 25926). For the main analysis, 53 participants had missing data and were excluded resulting in N=25,604 participants included in the linear models.

The significance of the associations between symptom scores and brain measures was determined using permutation testing, which accommodates potentially non-normally distributed data and covariance between predictor variables ^56^. To establish a permutation-based null-distribution, the analysis was repeated 5,000 times with the symptom severity scores randomly shuffled each time. The significance of the observed effect was then estimated by comparing the observed effect to the normally distributed, permutation-based null distribution using a two-sided t-test. An alpha level of 0.05 was used for all statistical tests and if applicable, correction for multiple testing was done through a false discovery rate-controlling procedure ^57^. The results section will explicitly state if the false discovery rate correction was applied, and the number of tests for which the correction was performed.

Subsequent analyses evaluated specificity of brain features for the severity of either insomnia, depression or anxiety symptoms. Bootstrapping was used to compare the strength of associations with the severity of each of the three symptom types. First, we generated a set of 5,000 bootstrap samples by randomly selecting participants from the original discovery dataset with replacement. Each bootstrap sample had the same number of participants as the original discovery dataset. Subsequently, we performed linear regression analysis on each bootstrap sample for each of the three symptom types. This procedure resulted in a distribution of 5,000 association strength values for each symptom. To test for symptom-specificity, indicated by significant differences in the association strength between the three symptom types, we conducted three two-sided t-tests: one between the insomnia and depression symptom association distributions, another between the insomnia and anxiety symptom association distributions and a third between the depression and anxiety symptom association distributions.

### Functional annotation: cognitive-emotional functioning

In order to evaluate *cognitive-emotional* functioning implications of possibly spatially distributed yet functionally related deviations, we performed enrichment analysis using the data-driven framework of neurobiological domains presented by Beam et al. ^58^. This framework identified brain regions associated with functional “domains” from human brain imaging literature. In our analyses, we used a division of the brain into eight domains that reflected distributed yet functionally integrated brain areas.

The volumetric division provided by Beam et al. ^58^ was mapped to the Desikan-Killiany atlas by calculating for each brain region its percentage overlap with each functional domain. We evaluated whether symptom-related deviations were enriched within any of these domains by computing for each functional domain the average association strength across brain regions weighted by the overlap of the regions with the domain. The significance of the average association of a functional domain with the severity of each symptom was assessed using permutation testing in which the linear regression was repeated 5,000 times with randomly shuffled symptom severities.

To evaluate the specificity of functional domains associated with the severity of each of the three symptoms, we first standardized the association between symptoms and brain measures across regions or connections. This standardization allowed us to compare the strength of associations between symptoms, regardless of potential differences in effect sizes resulting from different symptom severity assessment methods. The standardization process involved calculating the z-score of associations for each symptom and all brain regions or connections. The domain enrichment was then computed for the resulting standardized measures representing the aggregation of symptom-brain associations on a specific domain, referred to as the domain aggregation of symptom severity-brain associations. To identify differences in domain-aggregation across the three symptom types, we compared the aggregation across all possible symptom-pairs: insomnia and depressive symptoms, insomnia and anxiety symptoms and depression and anxiety symptoms. To assess the significance of these differences, we employed a bootstrapping procedure. This involved generating a set of 5,000 bootstrap samples by randomly selecting participants, with replacement, from the discovery dataset. Within each bootstrap sample, we performed linear regression analysis and calculated the domain-aggregation for the severity of each symptom type. To test the significance of the differences in domain-aggregation between two symptoms, we compared the associated domain-aggregation distributions using a two-sided t-test. This allowed us to determine if the domain-aggregation differed significantly between pairs of symptom types, thereby indicating symptom-specificity.

### Functional annotation: neurotransmission

To evaluate *neurotransmission* functioning implications, we examined the enrichment of symptom associations with respect to 19 different neurotransmitter receptors, receptor-binding sites and transporters. The data for this analysis were derived from PET tracer images of more than 1,200 healthy people presented in ^59^. We selected the top 25% regions with highest receptor and transporter densities as neurotransmitter-associated regions. The neurotransmitter association enrichment was then calculated and tested for significance using the methodology described above for cognitive-emotional domain-enrichment analysis.

### Functional annotation: large-scale functional resting-state networks

To evaluate functional network implications, we examined the enrichment of symptom associations with respect to seven functional networks: the visual, somatomotor, dorsal attention, ventral attention (also referred to as salience), limbic, frontoparietal (also referred to as central executive), and default mode network. The data for this analysis was obtained from the Yeo 7-network atlas that contains a parcellation of the 7 large-scale functional resting-state networks. An annotation file of the 7 functional networks was included for the fsaverage subject in the FreeSurfer Software package. We measured for each cortical region to what extent it was part of one of the 7 functional networks. For this, the surface-based annotation was translated to a 3D brain volume in volumetric space, in which each grey matter voxel was assigned a network label. Next, for each region in the Desikan-Killiany atlas, we computed the ratio of voxels that belonged to each of the 7 functional networks. We selected the top 15% regions that had the highest overlap with the functional networks as functional network-associated regions. The functional network association enrichment was then calculated and tested for significance using the methodology described above for functional annotation analyses. The functional network enrichment analysis is presented in the Supplementary Results.

### Potential confounding factors

We conducted a sensitivity analysis to assess the impact of potential confounding factors on our findings. We tested 10 possible confounders, detailed in the Supplemental Methods, as covariates in the analyses. Each of these factors was tested individually by repeating the analyses, with each iteration including one of the potential confounders as a covariate separately. To statistically determine if the results from this sensitivity analysis deviated from those of the main analysis, we tested whether the observed number of replicated regions was significantly lower than expected when compared to a reference distribution obtained using bootstrapping. Specifically, we repeated the analyses 5,000 times, each time randomly sampling participants, with replacement, from the main discovery sample to generate a distribution of the expected number of replicated regions.

### Sensitivity and validation analyses

Underscoring the potential influence of the used parcellation on brain measures, we repeated the analyses using a subparcellation of the Desikan–Killiany atlas ^60^ (DK-114, 114 cortical regions). The results are presented at the cognitive-emotional domain and neurotransmission system level in the Supplementary Results.

The spatial-anatomical specificity of the observed cognitive-emotional and neurotransmission system annotations was further assessed using a spin-permuted null model (10,000 permutations). We used the spin-permutation maps provided by Hansen et al. ^59^. These maps were created by randomly rotating the spherical projection of FreeSurfer’s fsaverage brain parcellation, with each region label being re-assigned the value of the nearest region label after rotation. This procedure was applied separately to each hemisphere. For subcortical areas, where spatial rotations are not feasible, regions were randomly shuffled in each permutation.

To evaluate the robustness of our reported results, we conducted multiple sensitivity analyses. In each sensitivity analysis, we tested the replicability of global brain measure findings and amygdala reactivity findings from the main analysis. Depending on the sample size, we either verified the replicability or examined the congruence of regional associations.

First, we assessed the robustness of our findings using imaging and questionnaire data obtained at a second imaging visit, which took place approximately 2-3 years after the first imaging visit and was available for N=3,601 participants. After MRI processing, quality control and checking for missing data the analysis sample was N=1,976. The symptom scores from the main analyses showed moderate to strong correlations with the scores obtained during the second imaging visit. Spearman’s rank correlation coefficients between the symptom severity measures were *ρ*=0.65 for insomnia symptoms, *ρ*=0.46 for depressive symptoms, and *ρ*=0.66 for anxiety symptoms. However, the small sample size of this second imaging visit sample limited the power to detect significant regional associations. Therefore, we quantified the generalizability of our results by computing the correlation coefficient between the regional associations observed in the main analysis and the regional associations observed in the second imaging visit. We interpreted the correlation coefficient by comparing it to the distribution of correlation coefficients that would be expected based on the results of the main analysis accounting for the smaller effect size. This distribution was estimated by bootstrapping a randomly selected sample of participants, with a similar number of participants as the in the second imaging visit dataset (N=1,976), from the discovery dataset 5,000 times.

Secondly, we assessed the generalizability of the results using a holdout sample of N=5,000 participants not included in the main analysis. After MRI processing, quality control and checking for missing data, the analysis sample was N=3,968. Similar to the sensitivity analysis using imaging data from the second imaging visit, we compared the results from this sample to the main findings and quantified them with respect to a bootstrapped sample of N=3,968 randomly selected participants from the main analysis.

Finally, to evaluate the impact of familial relationships on our findings, we performed a sensitivity analysis excluding 3,532 participants identified by the UK Biobank as first-degree relatives. As this analysis had a comparable number of participants as the main analysis, we tested generalizability of the results by the number of regional effects that could be replicated, similar to the methodology used in the sensitivity analysis with additional covariates.

## Data availability statement

The phenotype and MRI data from the UK Biobank that was used in this study can be accessed by researchers upon application, see:

https://www.ukbiobank.ac.uk/register-apply

Cognitive-emotional annotations were obtained using data available from:

https://github.com/ehbeam/neuro-knowledge-engine

Neurotransmission systems were derived from data available from:

https://github.com/netneurolab/hansen_receptors

## Code availability statement

Structural and functional connectivity was reconstructed using CATO, which source-code is available from:

https://github.com/dutchconnectomelab/CATO

MATLAB code used in the statistical analyses will be shared on GitHub:

https://github.com/Sleep-and-Cognition/multimodal-insomnia-depression-anxiety

## Acknowledgements

This work has received funding from ZonMw, the Hague, The Netherlands, project 09120011910032 REMOVE, the European Research Council (ERC), Brussels, Belgium, Advanced Grant 101055383 OVERNIGHT, the Dutch Research Council (NWO), the Hague, The Netherlands, VENI 201G.064 [to JES], and from the Dutch Research Council (NWO), the Hague, The Netherlands, VIDI 452-16-015 [to MPvdH], and from the ERC under the European Union’s Horizon 2020 research and innovation programme (Grant agreement No. ERC CONNECT 101001062) [to MPvdH].

