## Supplementary Materials for "Multimodal brain imaging of insomnia, depression and anxiety symptoms: transdiagnostic commonalities and differences"

### Supplementary Methods

*Quality control*

Besides filtering participants with MRI data that was indicated as unsuccessful by the UK Biobank or with errors in the processing pipeline, data driven quality control was applied. Imaging data were checked to ensure that T2-flair was used in addition to T1 to run FreeSurfer (field ID: 26500) and that the intensity scaling for T2-flair was larger than 3 (field ID: 25926), subjects had no missing longitudinal (z) brain position information (field ID: 25758), and no not-a-number items in the regional or connectivity measures, resulting in the exclusion of 184 subjects.

Further outliers were selected based on measures for MRI quality control provided by the UK Biobank ("discrepancy" between T1 image and template after linear alignment (field ID: 25731), "discrepancy" between T1 image and template after non-linear alignment (field ID: 25732), amount of nonlinear warping (field ID: 25733), discrepancy between T1 image and rs-fMRI after linear alignment (field ID: 25739), |discrepancy between T1 image and DTI after linear alignment (field ID: 25737), signal to noise ratio in rs-fMRI (field ID: 25744), rs-fMRI head motion (field ID: 25741), total number of outliers slices in DWI image (field ID: 25746), scanner lateral (X) brain position (field ID: 25756), scanner transverse (Y) brain position (field ID: 25757) and scanner longitudinal (Z) brain position (field ID: 25758). Individuals scoring 2.5 interquartile-range (IQR) below the first quartile (Q1) or above the third quartile (Q3) on any of these criteria were considered outliers.

Similar data-based outlier detection was used on the regional volume, surface and thickness measures, mean functional connectivity strength, amygdala responsivity and the average structural connectivity FA strength, the average group prevalence of structural connections present and the group-prevalence of absent connections. Subjects with any regional measure or connectivity summary measure that scored below Q1 - 2.5*IQR or above Q3 + 2.5*IQR on any of these measures were considered outliers. The quality control based on UK Biobank provided measures and data-based measures resulted in the exclusion of 6,092 subjects and a sample size of N=25,657.

*Insomnia symptoms*

The single UK Biobank question on insomnia has been validated in great depth by Hammerschlag et al.[1]. They demonstrated excellent sensitivity (98%) and specificity (96%) of the UKB question to distinguish cases, who met both the Insomnia Severity Index and Pittsburgh Sleep Quality Index criteria, from controls scoring below the cut-off scores. The UK Biobank item had similar excellent sensitivity (89%) and specificity (91%) to distinguish cases from controls who completed a structured diagnostic interview.

In sharp contrast to the validity of the single UK Biobank question, the use of a clinical diagnosis from medical records has paradoxically poor accuracy to distinguish cases from controls. The reason for this is that insomnia disorder is severely underreported and underdiagnosed in clinical practice (e.g. [2,3 4 5]) and alike in population based studies that make use of patient records. For example, in a sample of 42,507 patients, the diagnosis of insomnia as a primary or comorbid diagnosis was made only for 34 out (0.08%), even though about 40% of these patients experienced severe sleep disturbance and 22% reported this to be one of their most prominent problems [6].

*Depressive symptoms*

For depression, we utilized two questions that form the PHQ-2, a concise measure validated by Kroenke et al. [7]. This measure efficiently captures the core criteria for major depressive disorder diagnosis with good sensitivity (72%) and specificity (85%). Measuring the accuracy of the PHQ-2 in identifying participants with a history of having depression diagnoses (field ID: 20544, collected in 69% of the participants with processed MRI data), showed low sensitivity (42%) and good specificity (83%). We note that this might not justify the performance of the PHQ-2 questionnaire, because this compares the PHQ-2 to having a diagnosis at any point in the lives of the participants and it is expected that the performance would be better if the PHQ-2 was compared to having a recent (or current) depression diagnosis. More insightful might be comparing the PHQ-2 with the full PHQ-9 which is an often-used depression questionnaire [8]. The PHQ-2 focuses solely on depressed mood and loss of interest, thereby representing the DSM-IV diagnostic core criteria. Measuring the nonparametric correlation between the PHQ-2 and PHQ-9 for a subset (69%) of participants that had taken part in the optional "online mental health questionnaire" showed a strong correlation (Spearman's $\rho$=0.68, p<0.001).

*Anxiety symptoms*

Anxiety symptom severity was assessed using the anxious-tense factor [9,10] from the N-12 questionnaire [11], which aligns with previous research methodologies and is appropriate for the data available. In the UK Biobank sample, an anxious-tense score larger than zero showed 64% sensitivity and 67% specificity for self-reported lifetime "anxiety, nerves or generalized anxiety disorder" diagnosis (field ID: 20544). Similarly, this assessment might have been more accurate if tested against recent anxiety diagnoses. Among the subset of participants who engaged in the "online mental health questionnaire", the anxiety symptom score showed higher sensitivity (80%) and slightly lower specificity (63%) in classifying people that had severe anxiety according on the GAD-7 questionnaire [12] (cut-off >15 points, field IDs: 20505, 20506, 20509, 20512, 20515, 20516 and 20520).

*Alternative symptom scores*

For a subset of participants who had both in-depth phenotype data and quality-controlled MRI data available (N=15,357, 60% of the sample in the main analyses), we computed alternative anxiety and depression symptom severity scores for use in sensitivity analyses. For depressive symptoms, we combined the PHQ-2 score collected at two time points, the PHQ-9 score from a separate time point, and self-reported lifetime depression diagnosis (coded as zero or one). For anxiety symptoms, we combined the anxious-tense score from the N-12 questionnaire, the GAD-7 questionnaire score, and self-reported lifetime diagnoses of "anxiety, nerves, or generalized anxiety disorder" (also coded as zero or one). First, these measures were all standardized to have a mean of zero and a standard deviation of one. Then, for each symptom type, we computed the principal components and participants' scores on the first principal component were used as alternative anxiety and depression symptom severity scores.

*Potential confounding factors*

*Medication use.* We assessed medication use during the initial assessment center visit (field ID: 20003), coding it into four categories of psychiatric medications: anxiolytics, antidepressants, antipsychotics, and mood stabilizers.

*Body mass index (BMI).* BMI data were collected at the initial assessment by the UK Biobank (field ID: 21001). BMI is a recognized risk factor for the onset of mood and anxiety disorders [13] and has been associated with, partly methodological, differences in brain measures [14] [15].

*Social-economic status.* We used the Townsend deprivation index (Field ID: 22189) to assess socio-economic status [16]. This index is calculated by the UK Biobank using national census data, assigning scores based on the output area corresponding to each participant's postcode.

*Educational attainment.* We used years of education as measure for educational attainment and computed it using the same method as earlier studies [17,18]. The participants were grouped into 6 highest-received education qualifications (Field ID: 6138, qualifications included "CSEs or equivalent", "O levels/GCSEs or equivalent", "NVQ or HND or HNC or equivalent", "A levels/AS levels or equivalent", "College or University degree", "Other professional qualifications e.g.: nursing, teaching"). Participants that indicated no education ("None of the above") or zero years of education, were excluded. 65% of the participants had information available on the age that they completed full time education (Field ID 845). Missing values were imputed based on averages from similar educational levels and the estimated age of completing a college or university degree was assumed 22 years.

*Childhood stress.* Childhood stress was assessed by five questions that were modified versions of the Childhood Trauma Questionnaire [19,20] (Field IDs: 20487-20491). Responses were scored on a five-point scale, with reverse coding applied where applicable, and summed to create a childhood stress score.

*Adulthood stress.* Similar to childhood stress, adulthood stress was measured by five questions focusing on stressful experiences in adulthood, such as potentially abusive relationships (Field IDs: 20521-20525). These were also scored on a five-point scale with two items reverse coded and summed in an adulthood stress score.

*Smoking.* Smoking status (field ID: 20116) was dichotomized; individuals were coded 1 if they had ever smoked (in the past or currently) and 0 if they had never smoked.

*Diabetes.* Diabetes diagnostic status was derived from the health-related outcomes from the UK Biobank that were obtained through linkages to a range of health-related records including primary care data, hospital inpatient data, death register records and self-reported medical conditions. Diabetes was defined as either: “insulin-dependent diabetes mellitus”, “non-insulin-dependent diabetes mellitus”, “malnutrition-related diabetes mellitus”, “other specified diabetes mellitus” and “unspecified diabetes mellitus” (Field IDs: 130707, 130709, 130711, 130713, 130715, corresponding to e10, e11, e12, e13, e14).

*Hypertension.* Similar to the diabetes diagnostic data, hypertension was obtained from the health-related outcomes provided by the UK Biobank. Hypertension was defined as either "essential (primary) hypertension", "hypertensive heart disease", "hypertensive renal disease", "hypertensive heart and renal disease", "secondary hypertension" (Field IDs: 131287, 131289, 131291, 131293, 131295, corresponding to i10, i11, i12, i13, i15).

*Alcohol usage.* Alcohol consumption was quantified using a log-transformed measure of typical daily ethanol intake, as detailed in Thijssen et al. [21].

### Supplementary Results

*Functional annotation: Large-scale functional networks*

Functional annotation of our results with respect to seven large-scale functional networks revealed two symptom-specific patterns: Regions within the "limbic" network exhibited thinner cortices in proportion to the severity of depressive symptoms ($\beta$=-0.011, p=0.023), a finding significantly more pronounced than in insomnia ($p=0.030$) or anxiety ($p<0.001$, see Supplementary Figure 6). Additionally, weaker functional connectivity was observed in the frontoparietal network in relation to more severe insomnia ($\beta=-0.010$, $p=0.005$) and anxiety symptoms ($\beta=-0.012$, $p=0.005$), with these changes being significantly more pronounced than in depressive symptoms ($p=0.005$ and $p=0.027$, respectively).

*In-sample validation analysis*

To evaluate the robustness of our findings, we used quality-controlled imaging and questionnaire data of 1,976 participants that were reassessed during a second imaging visit approximately 2-3 years later. The three symptom severity scores showed moderate to strong correlations across the time-averaged scores used in the main analyses and those assessed during this second imaging visit: Spearman's rank correlation coefficients were $\rho=0.65$ for insomnia symptoms, $\rho=0.46$ for depressive symptoms and $\rho=0.66$ for anxiety symptoms.

The analysis of global brain measures in this sample showed partial replication, with 6 of the 11 initial findings being replicated (see Supplementary Table 5). Given the smaller sample size at the second imaging visit and the large number of tests performed in the regional analyses, our focus was on testing the alignment, rather than strict replication, of regional results from the main analysis and those of the second imaging visit. To assess consistency between the regional association maps of both analyses, we computed their correlation and compared the observed correlations against their expected ranges determined through bootstrapping. Instances with a lower-than-expected correlation, assessed by one-sided permutation testing, then indicated limited generalizability of findings. The correlation between the association strength maps generally fell within the expected range of correlations (see Supplementary Figure 4). However, three exceptions were noted where observed correlations were significantly lower than anticipated. These included the association of depression symptom severity with functional connectivity (with an $r_{observed}=-0.12$ and $r_{expected}=0.37$, $p=0.007$) and cortical thickness (with an $r_{observed}=0.17$ and $r_{expected}=0.52$, $p=0.027$), as well as for the initial main finding of associations of insomnia symptom severity with subcortical volumes (with an $r_{observed}=-0.28$ and $r_{expected}=0.52$, $p=0.009$).

*Out-of-sample validation analysis*

We assessed the generalizability of the results using a separate holdout sample that consisted, after quality control, of 3,986 participants who were not included in the discovery sample used for the main analyses.

In this holdout validation analysis, three of the 11 global brain measure findings were replicated (see Supplementary Table 5). In 9 of the 15 regional analyses (3 symptom types × 5 modalities), regional association maps exhibited patterns similar to the brain maps reported in the main analysis (see Supplementary Figure 5). Exceptions were the associations of regional cortical surface areas with severity of depressive symptoms, with an $r_{observed}$=0.05 and $r_{expected}$=0.52 ($p<0.001$), and anxiety symptoms, with an $r_{observed}$=0.08 and $r_{expected}$=0.49 (p=0.002). The correlation was also lower than expected for the regional associations of functional connectivity with insomnia symptom severity ($r_{observed}$=0.20 and $r_{expected}$=0.58, p=0.042). Finally, the correlation was lower than expected for the regional associations of structural connectivity with all symptom severities (insomnia: $r_{observed}$=-0.03, $r_{expected}$=0.45, p=0.003, depression: $r_{observed}$=0.00, $r_{expected}$=0.55, $p<0.001$, anxiety: $r_{observed}$=0.09, $r_{expected}$=0.47, p=0.003).

*Sensitivity analyses: familial relationships*

We assessed the effect of familial relationships in the UK Biobank dataset on our results. We excluded 3,532 participants that were identified by the UK Biobank as first-degree relatives of other participants to avoid potentially inflated significance levels and biases.

In this analysis, ten of the 11 initially reported global brain measure findings were replicated (see Supplementary Table 5). Given that the statistical power of this sensitivity analysis was comparable to the main study, we sought to ascertain the degree to which all regional associations identified in the primary analysis could be replicated. Across all imaging modalities, every regional association related to insomnia symptom severity was successfully replicated. Similarly, for associations with depressive symptom severity, findings were consistently replicated across all regions for the thickness, surface, volume, and functional connectivity modalities. In structural connectivity, the results aligned with the expected 95%-confidence interval, with 38 out of 42 regional effects being replicated (p=0.500). Regarding anxiety symptom severity, all regional effects were confirmed in the volume and functional connectivity modalities. Moreover, the number of replicated regions was within the 95%-confidence interval for the surface (58 out of 61 regional effects replicated, p=0.700) and structural connectivity (32 out of 38 regional effects replicated, p=0.600) modalities.

*Alternative Desikan-Killiany subdivision DK-114 atlas*

The DK114 atlas provided us the opportunity to validate our results with respect to a different brain parcellation. To compare the results between the main section and those obtained using this subdivision, we compared the functional annotation results that were defined for both atlases. See Supplementary Figure 7.

Focusing on symptom-specific results identified in the main analysis, we successfully replicated in the DK-114 atlas the depression-specific association of symptom severity with cortical thickness in regions linked to the "reward" and "language" domains. The symptom-specific association of both anxiety and depressive symptoms with cortical surface area in regions with high dopamine 2 receptor densities were only replicated for anxiety symptoms in the DK-114 atlas. The depression symptom-specific association of cortical thickness with the cannabinoid 1 neurotransmission system was not replicated.

The anxiety-specific functional annotations of functional connectivity with the H3, NMDA, mGluR5 and D1 receptors were replicated in the DK-114 atlas. The functional annotation of weaker structural connectivity with insomnia symptom severity in the “cognition” domain showed no symptom-specificity in the DK-114 atlas, but the functional annotation itself remained significant.

*Spin-based permutation testing*

The spatial-anatomical specificity of the observed cognitive-emotional and neurotransmission system annotations was further tested using a spin-permuted null model [22]. In brief, this method compares the observed association between two brain maps with a null-model obtained by randomly rotating the examined brain maps on a spherical projection of the brain.

Our results showed that the significant associations identified with structural and functional connectivity were significant under spin testing indicating functional specificity (see Supplementary Figure 8). However, the functional annotation of associations between the symptom severity scores and cortical surface area, cortical thickness and subcortical volume were only partly replicated, suggesting these findings might be due to brain-wide cortical surface area changes and general subcortical effects rather than being specific to cognitive-emotional or neurotransmission systems.

*Alternative symptom scores*
The alternative depression and anxiety symptom severity scores showed somewhat higher intercorrelations compared to the scores used in the main analyses (Spearman’s rank correlation coefficients: insomnia-depression $\rho$=0.27, p<0.001; depression-anxiety $\rho$=0.46, p<0.001; anxiety-insomnia $\rho$=0.20, p<0.001, all FDR-corrected).

In all 10 regional analyses (2 symptom types × 5 modalities), the regional association maps derived using the alternative symptom severity scores correlated well with those from the main analysis, not falling below the 95%-confidence interval of the expected correlation as determined by bootstrapping (see Supplementary Figure 9). This indicates that the alternative symptom severity scores show brain property associations consistent with the findings from the main analyses.

Next, we looked at the number of regions that were reported significant in the main analyses and could be validated using the alternative symptom scores. In 9 out of 10 regional analyses, the number of confirmed regional associations was within the expected 95%-confidence interval obtained using bootstrapping. The exception was the association between anxiety symptoms and cortical surface area, where only 23 regions replicated out of 61 regions reported significant in the main analyses (expected number was 47 regions, 95%-confidence interval = [26-61], p=0.040). Notably, the expected range of the number of replicated regions was generally broad across modalities. For example, the replication of zero regional associations between depressive symptom severity and subcortical volumes was still within the expected range. This suggests that this sensitivity analysis was limited in statistical power, due to the limited number of participants (60%) that had in-depth symptom and diagnosis data available.

**Supplementary discussion**

*Cross-modal integration*

The integrated analysis of multiple modalities enabled us to examine if significant correlations in symptom-related brain deviations occurred across modalities. By correlating regional symptom-related deviations across modalities post-hoc analysis, we identified modalities with convergent or divergent deviations (see Supplementary Figure 10).

For insomnia symptom severity, we found significant positive correlations between the regional deviations in cortical surface area and those seen in structural and functional connectivity ($r=0.40$, $p=0.018$ and $r=0.40$, $p=0.048$, respectively, all FDR-corrected for multiple testing). Looking at the cognitive-emotional domain annotations, we noticed that deviations centered around the “cognition” domain, which showed significant enhancement across these modalities.

In contrast to the other symptom types, we found for insomnia symptom severity a (non-significant) negative correlation between subcortical volume and functional connectivity ($r=-0.32$, $p=0.362$) between the regional brain correlates of these modalities. Further exploring this potential relation, we observed that while the amygdala showed significantly weaker functional connectivity in individuals with more severe insomnia symptoms, its volumetric properties were not significantly different. This suggests a possible dissociation between structural changes and functional alterations in this region in people experiencing insomnia symptoms.

Depressive symptom severity was associated with a thicker cortex specifically in areas where there was a reduction in surface area ($r=-0.48$, $p<0.001$), notably within the occipital lobe which is crucial for visual processing. A positive correlation was observed between depressive symptom related cortical surface area and structural connectivity associations ($r=0.39$, $p=0.016$). Inspecting the regions and cognitive-emotional domains associated with depressive symptom severity in these modalities did not show any specific regions or domains that drove this effect.

The regional associations with anxiety symptom severity showed no significant correlations across modalities, although we observed high, non-significant correlations between smaller subcortical volumes and weaker structural and functional connectivity. In these modalities, we had observed prominent changes in the amygdala suggesting that amygdala disruption might be a common substrate for anxiety-related alterations across modalities.

*Neurotransmission systems*

The functional annotation of the symptom associations with respect to receptor and neurotransmitter systems provides valuable insights that could, potentially, inform therapeutic drug targets.

In our examination, we found that thinner cortices in proportion to depressive symptom severity mapped on regions with higher cannabinoid CB1 receptor density, and significantly more so than was the case for the severity of insomnia and anxiety symptoms. However, this effect was not replicated in the more fine-grained DK-114 atlas. Previous studies have linked cortical thinning of regions rich in CB1 receptors to cannabis use in people with schizophrenia [23], suggesting that investigating substance use in individuals with depression symptoms might be a valuable direction for future studies.

Anxiety symptom severity was, in comparison to depressive or insomnia symptoms, notably associated with weaker connectivity between regions with high densities of dopamine D1, histamine H3, NMDA, and metabotropic glutamate 5 (mGluR5) receptors. These results align with prior studies implicating H3 and mGluR5 in the pathophysiology of (chronic) anxiety in mice [24,25], and earlier research linking D1 dopamine binding to amygdala reactivity in response to fearful faces [26].

More specific to the severity of depressive and anxiety symptoms than insomnia symptoms were smaller cortical surface areas in regions enriched with dopamine D2 receptors. This supports earlier studies showing that genetic variability in dopamine D2 receptor variants correlates with sociability which is also genetically linked to depression and social anxiety [27].

Regarding insomnia and anxiety symptoms, we observed weaker connectivity, in comparison to depressive symptoms between regions enriched with α4β2 acetylcholine receptor densities. Nicotine, primarily obtained from cigarette smoking, interacts primarily with the α4β2 receptor and is associated with disrupted sleep [28]. Further, the α4β2 has been implicated in the anxiety-relieving effects of nicotine [29]. Our analysis suggests that tobacco use could be a relevant factor in insomnia and anxiety symptoms for future research. Importantly, including smoking as a covariate did not change these outcomes, indicating that while nicotine influences α4β2 receptor functioning, the observed changes in connectivity might not be directly due to smoking behavior.

In interpreting these findings, it is crucial to consider that the receptor density maps used were limited to cortical areas, potentially overlooking significant receptor densities in subcortical regions. Moreover, the receptor maps provide only a spatial distribution of the receptors involved and spatial co-occurrence of receptors and symptom related brain deviations do not necessarily imply interactions between them. Additionally, neurotransmission systems are complex and likely associated with insomnia, depression, and anxiety through dynamics that extend beyond the spatial co-occurrence reported here. For instance, although the histamine H3 receptor is targeted for sleep disorders [30], we did not find insomnia-specific clustering of brain deviations around the H3 receptor.

### Supplementary Figures

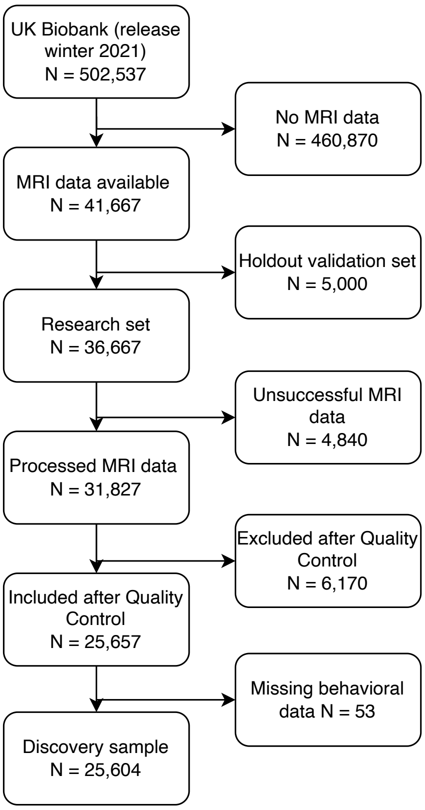


**Supplementary Figure 1. Flow diagram of UK Biobank participants included in this study.**

*The flow diagram shows the process of selecting UK Biobank participants for the current analysis. From the total number (N=502,537), MRI data were available for N=41,667 participants, of which 5,000 participants were randomly selected as holdout validation dataset. Of the resulting N=36,667 participants, 4,840 participants had unsuccessful MRI data, another 6,170 were discarded after quality control, and 53 participants had missing behavioral data resulting in N=25,604 participants included in the discovery sample.*

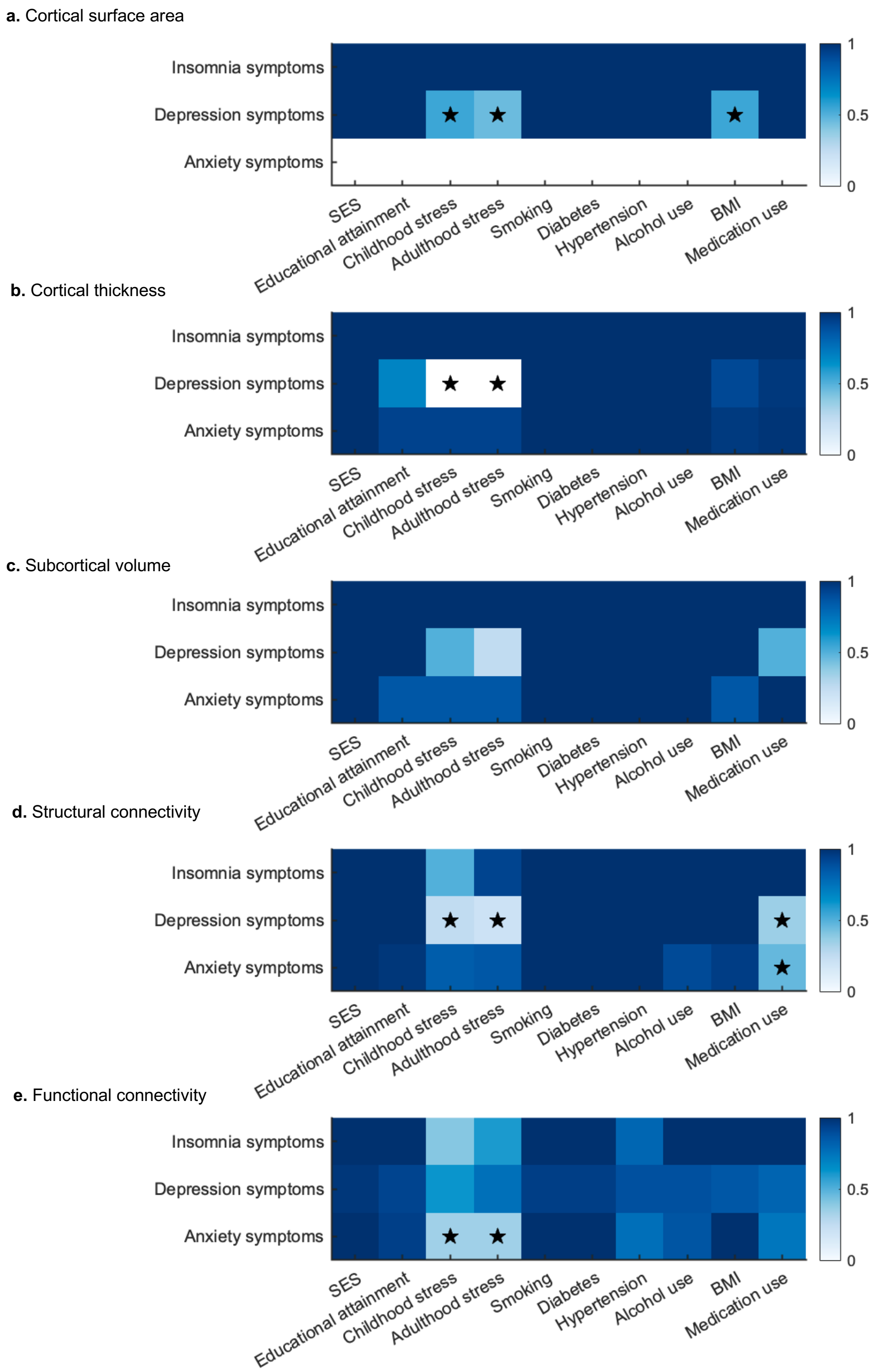


***Supplementary Figure 2. Potential confounding factors.*** *Ratio of regions that were replicated from the main analysis per symptom type and brain modality when additional covariates were included. Brain associations where less than expected brain regions were replicated (p<0.05), are indicated by a star (✶).*

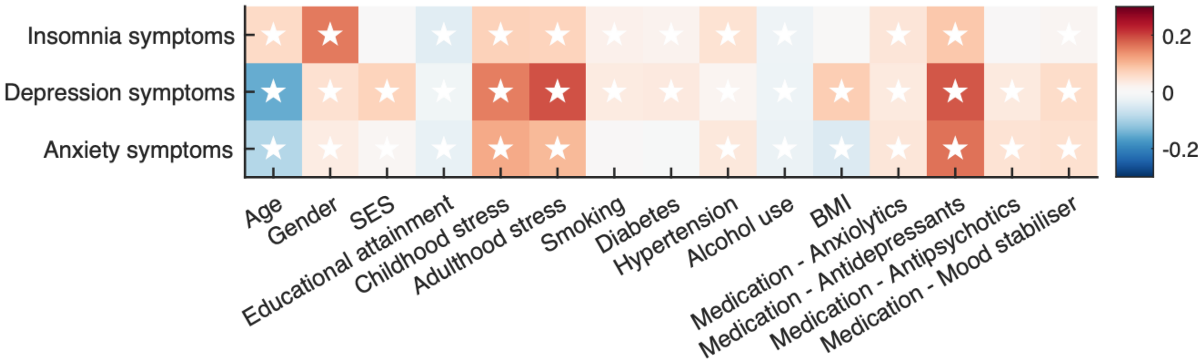


***Supplementary Figure 3. Correlations across symptom severity scores and potential confounding factors.*** *Spearman's rank correlation between the three symptom types and the 10 examined potential confounders. Medication use is subdivided into the 4 examined medication categories. Significant effects ($p<0.05$, FDR-corrected across 39 tests) are indicated by a white star (✶).*

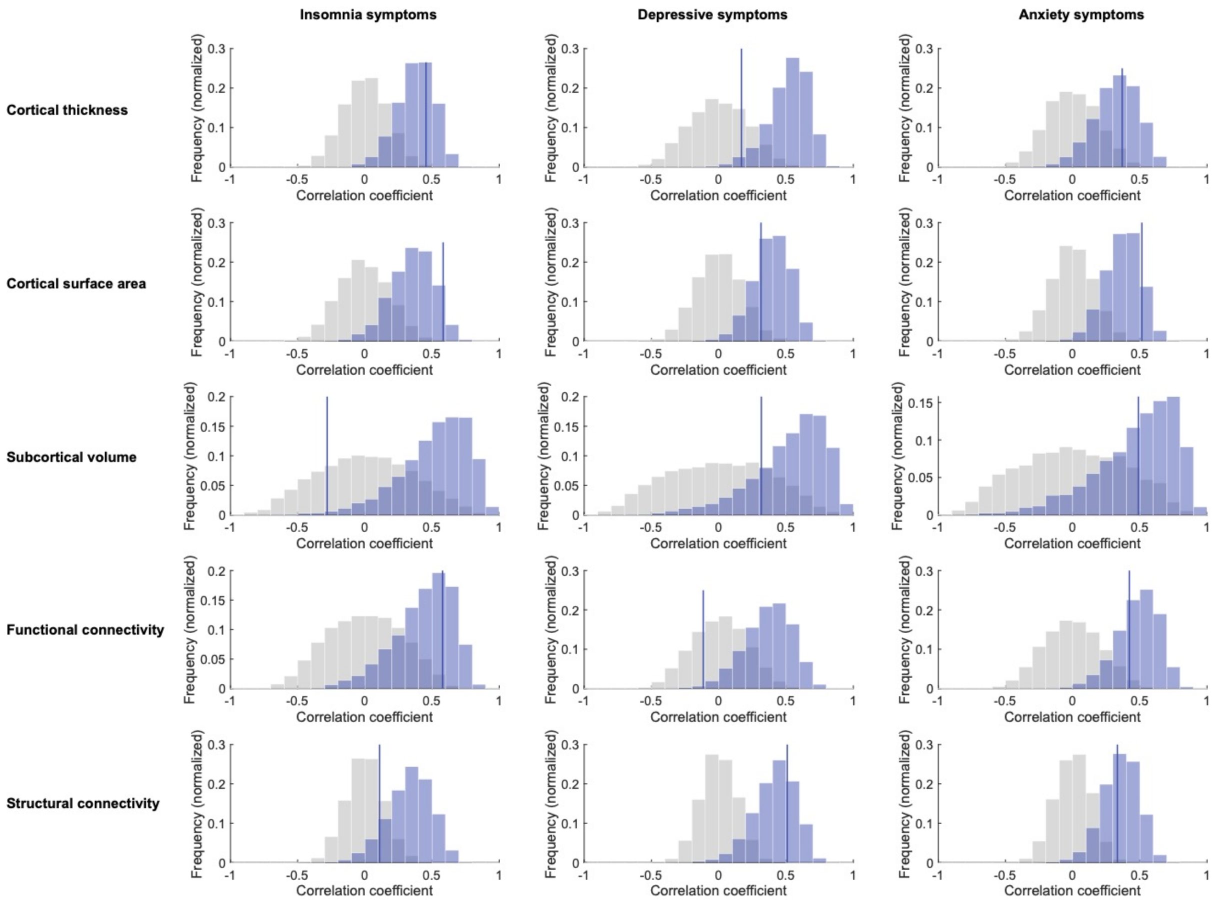


***Supplementary Figure 4. Correlation of regional brain-symptom associations obtained in the main analysis and in the second imaging visit dataset.*** *Dark blue lines indicate the observed correlation between the two datasets with respect to the profile of regional associations with symptom severity. Blue bars indicate the correlation coefficient distribution that would be expected based on the data from the main analyses (obtained using bootstrapping with 5,000 samples). Gray bars indicate the correlation coefficient distribution that would be expected if no brain-symptom associations were present (based on 5,000 permutations of the data from the second imaging dataset in which the symptom scores were shuffled across participants).*

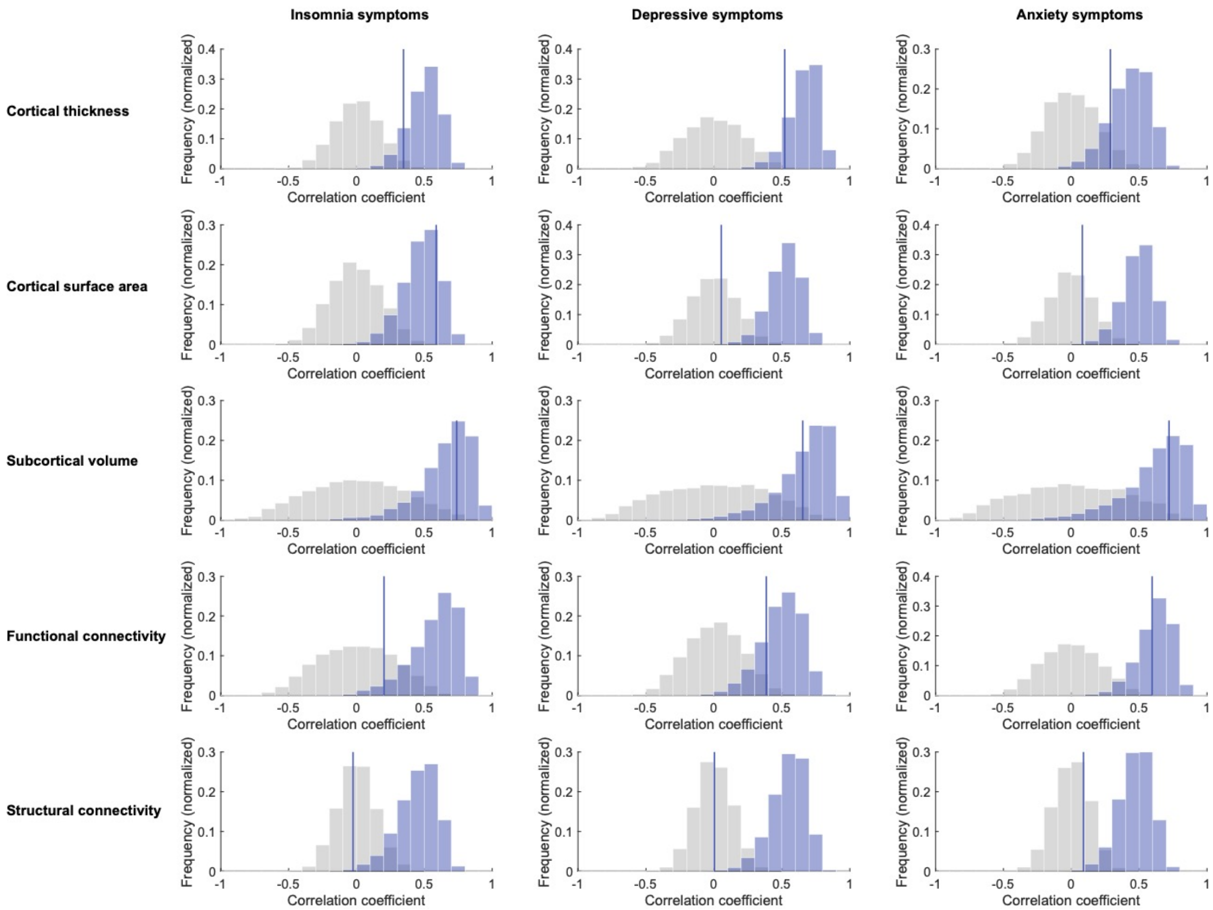


***Supplementary Figure 5. Correlation of regional brain-symptom associations obtained in the main analysis and in the holdout dataset.*** *Dark blue line indicates the observed correlation between the two datasets with respect to the profile of regional associations with symptom severity. Blue bars indicate the correlation coefficient distribution that would be expected based on the data from the main analyses (obtained using bootstrapping with 5,000 samples). Gray bars indicate the correlation coefficient distribution that would be expected if no brain-symptom associations were present (based on 5,000 permutations of the data from the holdout dataset in which the symptom scores were shuffled across participants).*

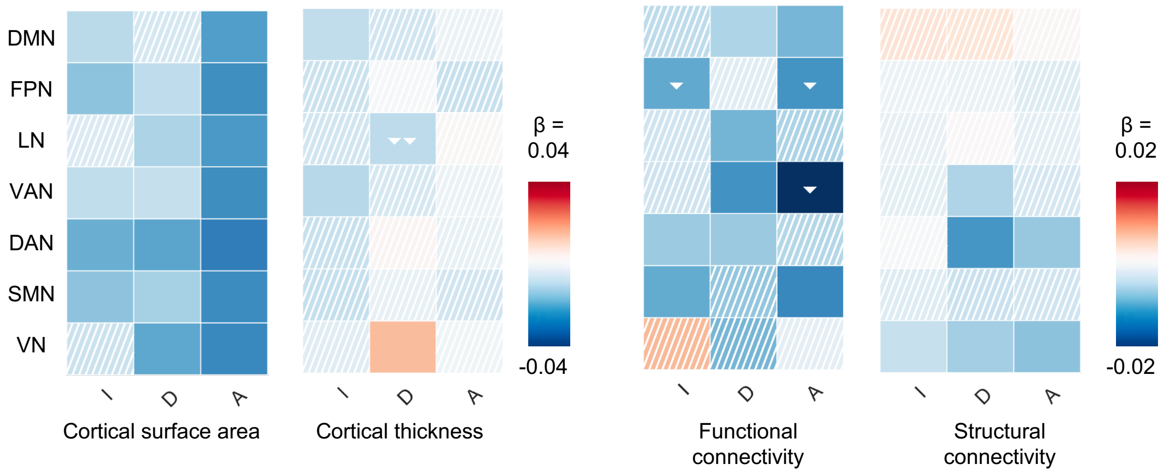


***Supplementary Figure 6. Large-scale resting-state functional networks annotation.*** *Enrichment analysis evaluated whether regional symptom-related variations were enriched within any of the 7 large-scale cortical resting-state functional networks identified by Yeo et al. [31], including the visual (VN), somatomotor (SMN), dorsal attention (DAN), ventral attention (VAN), limbic, frontoparietal (FPN), and default mode network (DMN). The correlation maps show the associations between symptom severities and the cortical surface area and thickness, and cortical functional and structural connectivity of regions assigned to the functional networks. Associations that are statistically significant (p<0.05 after FDR-correction across 21 tests [3 comparisons x 7 networks]) are depicted in solid color, while non-significant associations are marked with hatching. Negative effects (in blue) denote that the involved regions had smaller brain measures in specific functional networks. Additionally, white arrows highlight that, compared to another symptom, the association of the severity of a symptom is significantly more aggregated in areas involved in the indicated domain. Two white arrows in one cell indicate that the symptom severity-cortical thickness associations were more aggregated in areas involved in the indicated domain relative to both other symptom types.*

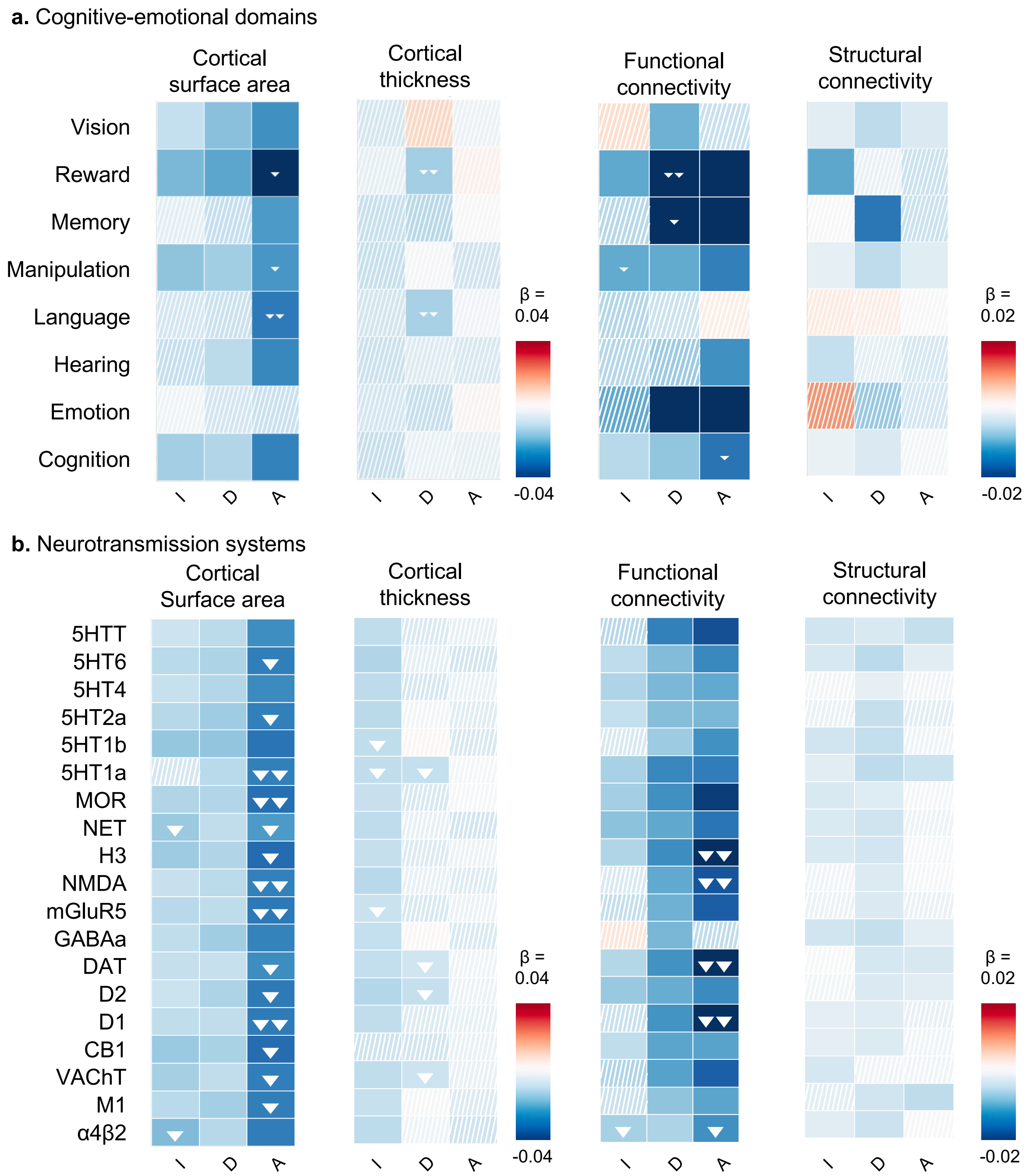


***Supplementary Figure 7. Functional annotation results obtained using the DK-114 cortical atlas.*** *Enrichment analysis evaluated whether regional symptom-related variations were enriched within any cognitive-emotional or neurotransmission system.* ***a.*** *These correlation maps show the associations between symptom severities and the cortical surface area, thickness in regions, and structural and functional connectivity between regions linked to cognitive-emotional domains. Associations that are statistically significant (p<0.05 after FDR-correction across 24 tests for cortical surface and cortical thickness [3 comparisons x 8 domains]) are*

*depicted in solid color, while non-significant associations are marked with hatching. Negative effects (in blue) denote that the involved regions had smaller surface area, thickness or volume in specific cognitive-emotional domains or neurotransmission systems. Additionally, white arrows highlight that, compared to another symptom, the association of the severity of a symptom is significantly more aggregated in areas involved in the indicated domain.* ***b.*** *Correlation maps that present the associations of symptom severities with brain measures of the top 25% cortical regions that have the highest densities for each of 18 receptor and transporter types (p-values are FDR-corrected across 54 tests [3 comparisons x 18 receptors and transporters]).*

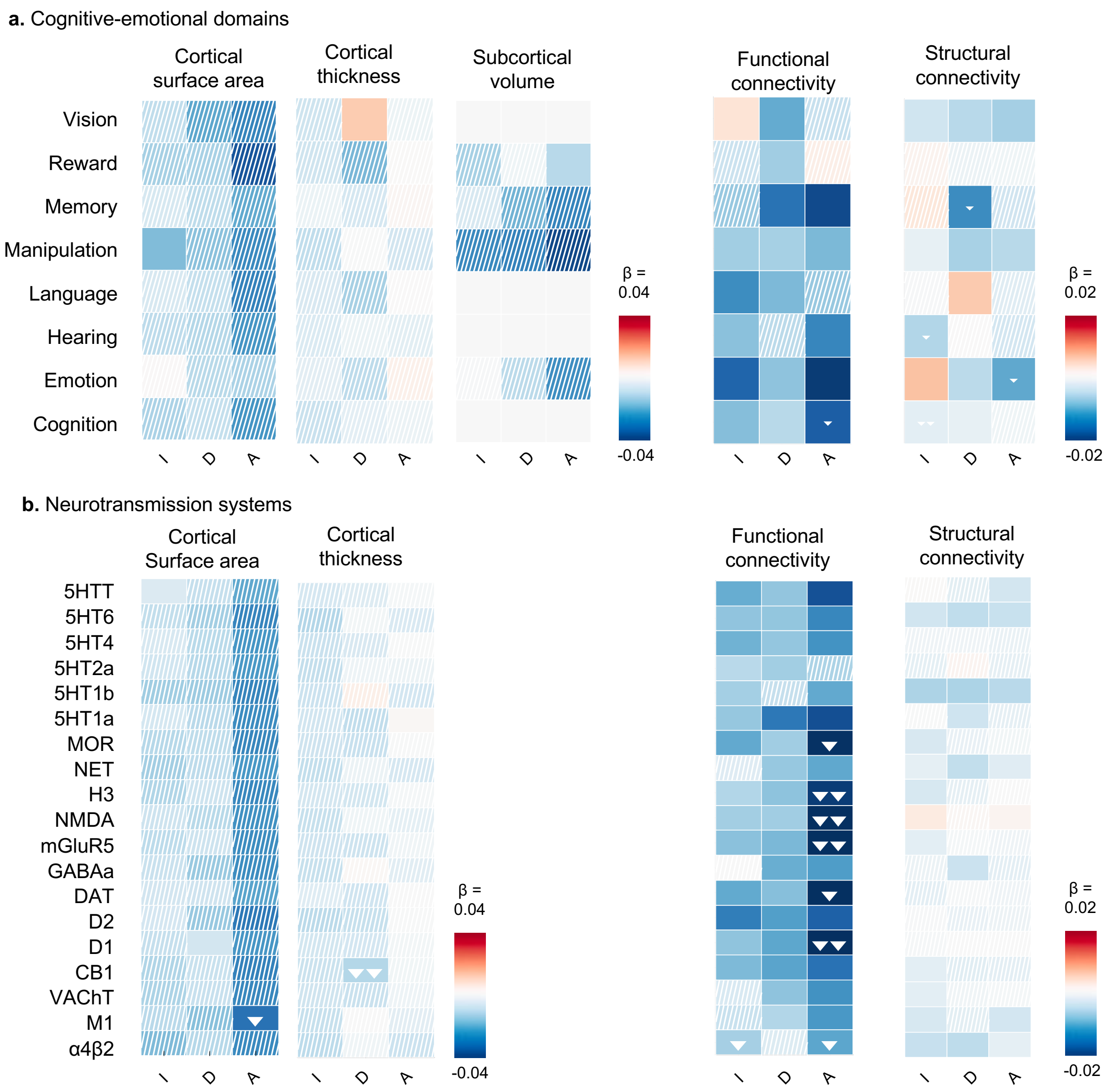


***Supplementary Figure 8. Functional annotation results obtained using spin-based spatial permutation testing.*** *Enrichment analysis evaluated whether regional symptom-related variations were enriched within any cognitive-emotional or neurotransmission system.* ***a.*** *These correlation maps show the associations between symptom severities and the cortical surface area, thickness and subcortical volume of regions, and structural and functional connectivity between regions linked to cognitive-emotional domains. Associations that are statistically significant (p<0.05 after FDR-correction across 24 tests for cortical surface and cortical thickness [3 comparisons x 8 domains], and 12 tests for subcortical volume [3 comparisons x 4 domains]) are depicted in solid color, while non-significant associations are marked with hatching. Negative effects (in blue) denote that the involved regions had smaller surface area, thickness or volume in specific cognitive-emotional domains or*

*neurotransmission systems. Additionally, white arrows highlight that, compared to another symptom, the association of the severity of a symptom is significantly more aggregated in areas involved in the indicated domain.* ***b.*** *Correlation maps that present the associations of symptom severities with brain measures of the top 25% cortical regions that have the highest densities for each of 18 receptor and transporter types (p-values are FDR-corrected across 54 tests [3 comparisons x 18 receptors and transporters]).*

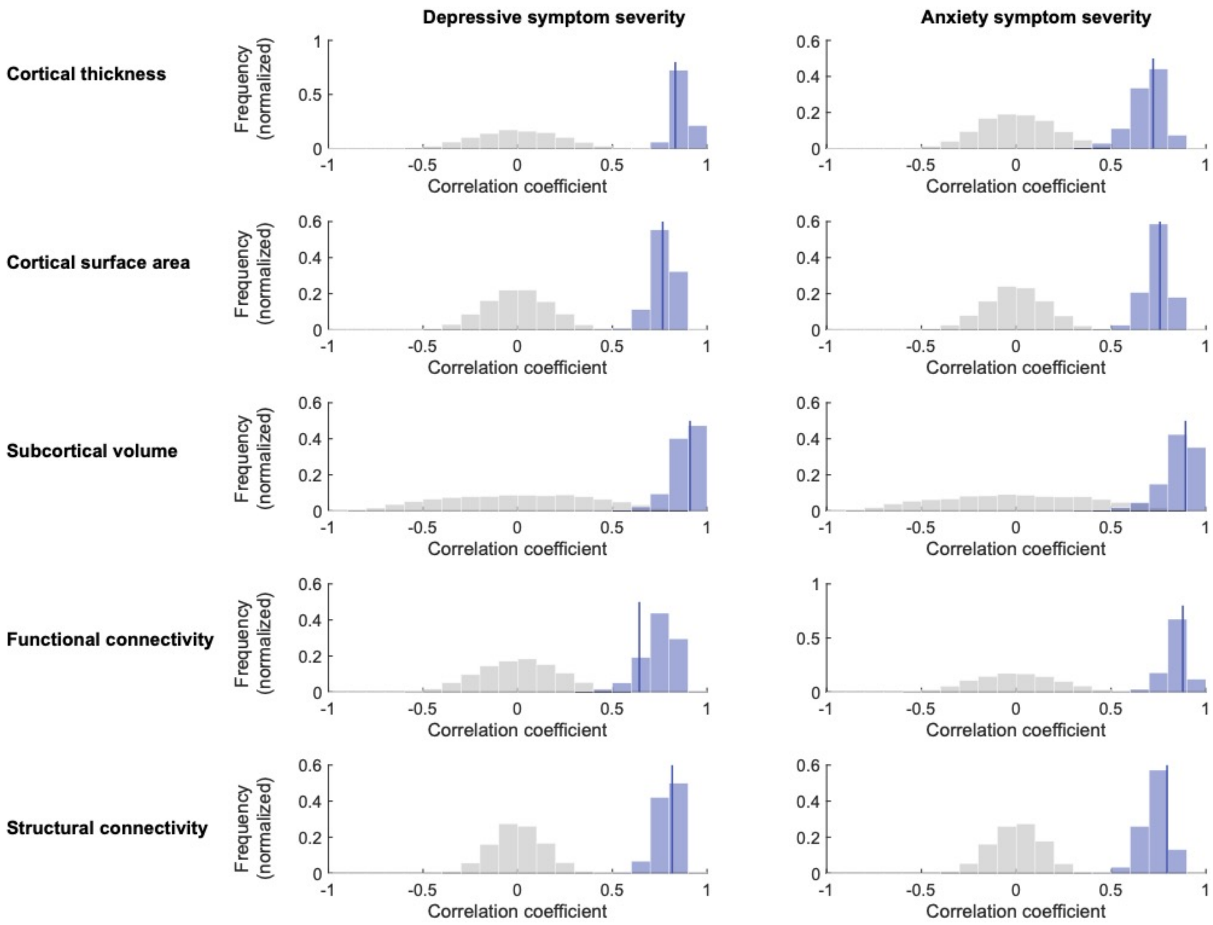


***Supplementary Figure 9. Correlation of regional brain-symptom associations obtained in the main analysis and using alternative symptom severity scores.*** *Dark blue lines indicate the observed correlation between the regional association maps obtained using the two symptom severity definitions. Blue bars indicate the correlation coefficient distribution that would be expected based on the data from the main analyses (obtained using bootstrapping with 5,000 samples with N=15,357). Gray bars indicate the correlation coefficient distribution that would be expected if no brain-symptom associations were present (based on 5,000 permutations).*

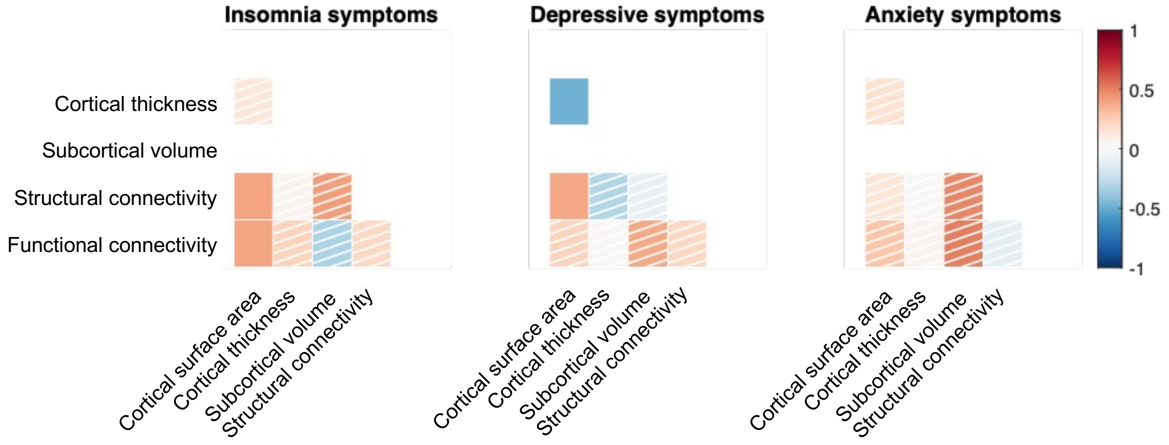


***Supplementary Figure 10. Correlation of Brain Correlates Across Modalities.*** *Correlation analysis of regional brain associations across different modalities revealed distinct correlation patterns for the three symptom types. Associations that are statistically significant (p<0.05 after FDR-correction across 8 tests) are depicted in solid color, while non-significant associations are marked with hatching.*

**Supplementary Tables**

*Supplementary* Tables *1-4 are in separate files.*

| Modality | Analysis | I | D | A |
|---|---|---|---|---|
| **Total cortical surface area** | *In-sample validation analysis* | β=-0.055, **p=0.042** | β=-0.063, **p=0.024** | β=-0.080, **p=0.004** |
| | *Out-of-sample validation analysis* | β=-0.033, p=0.099 | β=-0.006, p=0.768 | β=-0.052, **p=0.009** |
| **Average cortical thickness** | *In-sample validation analysis* | β=-0.021, p=0.379 | NA | NA |
| | *Out-of-sample validation analysis* | β=0.021, p=0.223 | NA | NA |
| **Subcortical volume** | *In-sample validation analysis* | β=-0.051, p=0.068 | NA | β=-0.063, **p=0.023** |
| | *Out-of-sample validation analysis* | β=0.010, p=0.598 | NA | β=-0.026, p=0.160 |
| **Average structural connectivity strength** | *In-sample validation analysis* | NA | β=-0.051, **p=0.023** | β=-0.045, p=0.052 |
| | *Out-of-sample validation analysis* | NA | β=-0.003, p=0.877 | β=-0.003, p=0.859 |
| **Average functional connectivity strength** | *In-sample validation analysis* | β=-0.041, p=0.069 | β=-0.007, p=0.786 | β=-0.052, **p=0.020** |
| | *Out-of-sample validation analysis* | β=-0.004, p=0.786 | β=-0.064, **p<0.001** | β=-0.038, **p=0.020** |

***Supplementary Table 5. Replication outcomes for global brain measures linked to symptoms of insomnia (I), depression (D), and anxiety (A).*** *The main analysis identified 11 significant associations between the global brain measures and the three symptom types. Each sensitivity and validation analysis aimed to replicate these findings. Successful replications with significant effects (p<0.05) are highlighted in bold.*
